# Mechanically Tunable, Compostable, Healable and Scalable Engineered Living Materials

**DOI:** 10.1101/2024.01.31.578234

**Authors:** Avinash Manjula-Basavanna, Anna M. Duraj-Thatte, Neel S. Joshi

## Abstract

Novel design strategies are essential to realize the full potential of Engineered Living Materials (ELMs), including their biodegradability, manufacturability, sustainability, and ability to tailor functional properties. Toward these goals, we present **Me**chanically **E**ngineered Living Material with **C**ompostability, **H**ealability, and **S**calability (**MECHS**) – a material that integrates these features in the form of a stretchable plastic that is simultaneously flushable, compostable, and exhibits the characteristics of paper. This plastic/paper-like material is produced directly from cultured bacterial biomass (40%) producing engineered curli protein nanofibers in scalable quantities (0.5-1 g L^-1^). The elongation at break (1-160%) and Young’s modulus (6-450 MPa) of MECHS was tuned to more than two orders of magnitude. By genetically encoded covalent crosslinking of curli nanofibers, we increase the Young’s modulus by two times. MECHS biodegrades completely in 15-75 days, while its mechanical properties are comparable to petrochemical plastics and thus may find use as compostable materials for primary packaging.

## Introduction

The emerging field of ELMs employs synthetic biology design principles to harness the programmability and the manufacturing capabilities of living cells and produce functional materials. ^1-4^ ELMs research not only provides avenues to integrate life-like properties into materials but also aims to realize *de novo* functionalities that are not found in natural or synthetic materials. ^5-21^ In recent years, several ELMs have been developed to demonstrate various functionalities such as adhesion, catalysis, mineralization, remediation, wound healing, and therapeutics *etc*. ^22-31^ ELMs that are mechanically stiff or soft have also been reported but the rational modulation of mechanical properties to a wide range through genetic programming remains elusive. ^5,6,9-11,25,32^ In this regard, we present a new ELM called MECHS that is fabricated at ambient conditions by a new method that comprises of genetic encoding, tailoring of mechanical properties, scalable production, healability and compostability (**Figure 1**).

**Figure 1:**
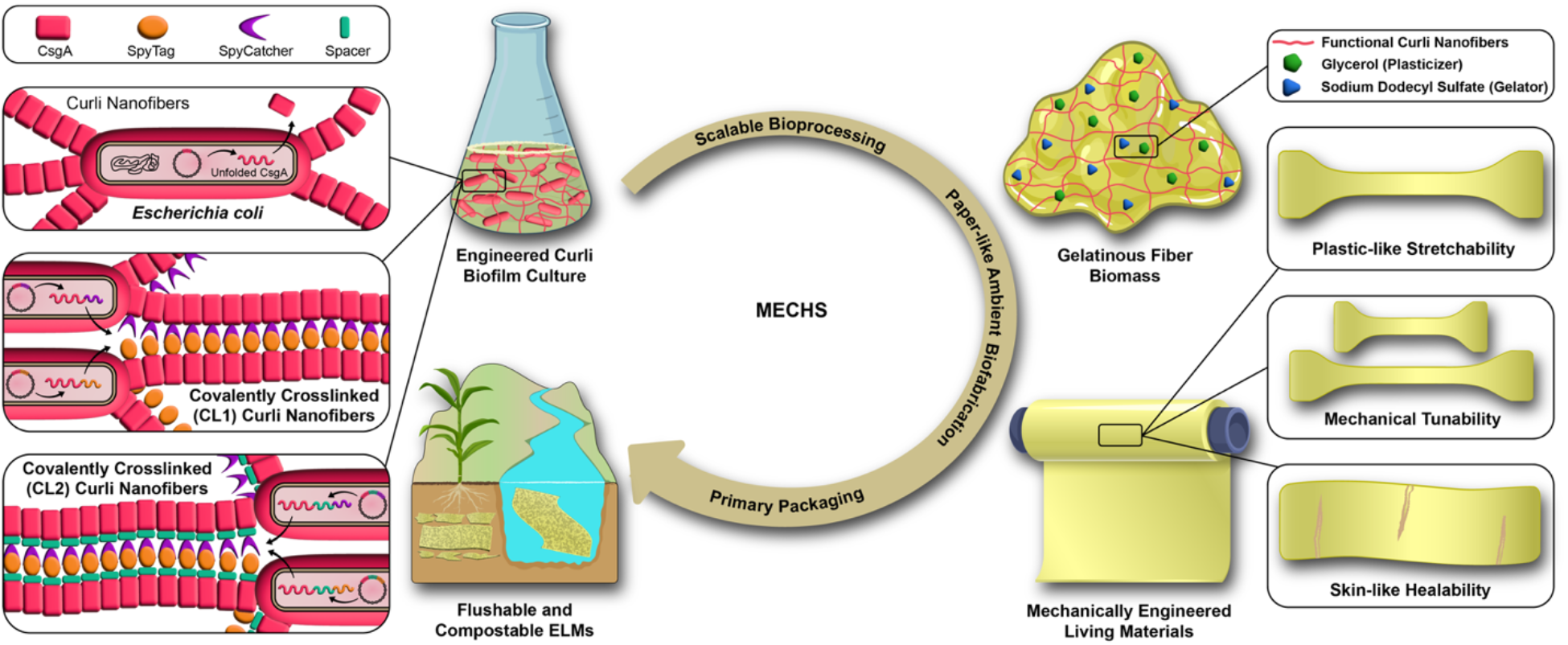
Schematic summary of Mechanically Engineered Living Materials with Compostability, Healability and Scalability (MECHS). Native and functional curli nanofibers were produced from engineered *Escherichia coli* and the treated biomass was dried ambiently to biofabricate MECHS films in a scalable manner. MECHS films exhibit plastic-like stretchability, mechanical tunability and skin-like healability. Parts of the schematics were adapted from BioRender.com.

Advances in biomanufacturing are extremely important at a time when human-made materials have been estimated to outweigh all the living biomass of planet Earth. ^33^ The existing paradigm of a linear materials economy (make-use-dispose) for synthetic materials is causing potentially irreversible damage to our ecosystem in the form of pollution and global warming. While many strategies will need to be employed to address these challenges, it is clear that bio-based manufacturing will need to be part of the solution. ^34^ Inspired by natural systems that utilize sustainable feedstocks and energy-efficient processes, coupled with their biodegradation to initiate a new cycle, biomanufacturing should strive to create materials that have similar recyclability or potential for conversion to benign components to create a circular material economy. ^35-37^ Such nature-inspired sustainable solutions enabled by biomanufacturing will also make inroads toward practical implementation through a combination of appropriate government policies, public interest, and investment. ^38^

Previously, we had reported a new bioplastic known as AquaPlastic composed of recombinant protein nanofibers produced by *E. coli*. ^9^ It exhibited a Young’s modulus of ∼1 GPa and ultimate tensile strength of ∼25 MPa, comparable to petrochemical plastics and other bioplastics. ^9^ AquaPlastic was also resistant to various chemicals (*e*.*g*., acid, base, and organic solvents), and could adhere to and coat a wide range of surfaces, protecting them from wear and environmental conditions. ^9^ However, the broad utility of AquaPlastic was limited due to its brittleness and lack of scalability. Additionally, we had earlier shown that whole microbial biomass could be dried to form cohesive and glassy stiff materials with a streamlined fabrication and higher yields compared to AquaPlastic, at the expense of tunability. ^12^ Here we report a new fabrication strategy to combine whole cellular biomass and engineered extracellular matrix protein nanofibers that enables tuning of their mechanical properties. Our new material, MECHS, exhibits properties similar to both plastic and paper, showcasing: 1) a new fabrication strategy that enables larger scale production of flexible films at ambient conditions, analogous to paper manufacturing; 2) genetic engineering to tailor their tensile stiffness and strength; 3) compositional and morphological analysis; 4) compostability, 5) a landscape of achievable mechanical properties comparable to conventional petrochemical plastics, bioplastics and other relevant bio-and synthetic materials; and, 6) prototypes for disposable packaging applications, contributing to the creation of a sustainable circular material economy.

### Biofabrication of MECHS

MECHS is fabricated from a combination of whole *E. coli* cells and engineered recombinant curli nanofibers. Curli are an extracellular matrix component of microbial biofilms and are composed of nanofibers self-assembled from a protein building block, CsgA (**Figure 1**). ^39^ Curli nanofibers are resistant to heat, solvents, pH, detergents, and denaturants, and thus serve as a good biopolymeric scaffold for robust materials. ^40^ To express the recombinant curli nanofibers, we used an *E. coli* strain that we previously developed (PQN4), in which the chromosomal curli genes (*csgBAC, csgDEFG*) have been deleted. ^41^ PQN4 was transformed with a pET21d plasmid vector encoding a synthetic curli operon, *csgBACEFG*, containing all the genes necessary for CsgA production, secretion, and extracellular assembly. In a typical biofabrication of MECHS, the curli-containing *E. coli* biomass was treated with 1-5% (w v^-1^) of sodium dodecyl sulfate (SDS) to obtain a gelatinous substance, which enables facile casting in a silicone mold. Ambient drying in the mold resulted in films that were brittle and, in some cases, (1% and 2% SDS) convoluted (**Figure S1, S2**). To achieve flexible MECHS films, we added glycerol (1-5% w v^-1^), a plasticizer commonly used with bioplastics, to the gelatinous curli biomass prior to casting (**Figure S3, S4**). ^42^

MECHS films that had been pre-treated only with SDS (i.e., “gelator”) and no glycerol (i.e., “plasticizer”), were extremely brittle as measured by tensile mechanical tests, with elongation at break values of 0.6 ± 0.4% (**Figure 2a-f, S5a** and **S6**). With 1% plasticizer, the elongation at break was found to increase considerably to 10.2 ± 6.9% (**Figure 2c-g** and **S5b**). Similarly, as the plasticizer content increased to 2%, 3%, 4% and 5%, the elongation at break increased significantly to 35.5 ± 7.7%, 70.1 ± 16.3%, 101.9 ± 28.8%, and 159.3 ± 25% respectively (**Figure 2c-f,h-k** and **S5c-f**). On the other hand, the corresponding Young’s modulus decreased from 450 ± 206.4 MPa to 6.6 ± 1.7 MPa as the plasticizer amount increased (**Figure 2e**). Ultimate tensile strength values of MECHS films also decreased with increasing plasticizer (**Figure 2f**). Overall, our method further streamlines the fabrication of flexible MECHS films from our previous demonstrations by casting directly from whole microbial biomass, without the need for filtration and extensive washing. ^9^ However, it also provides an opportunity to tailor their mechanical properties by two orders of magnitude by inclusion of the engineered curli nanofibers and a plasticizer.

**Figure 2:**
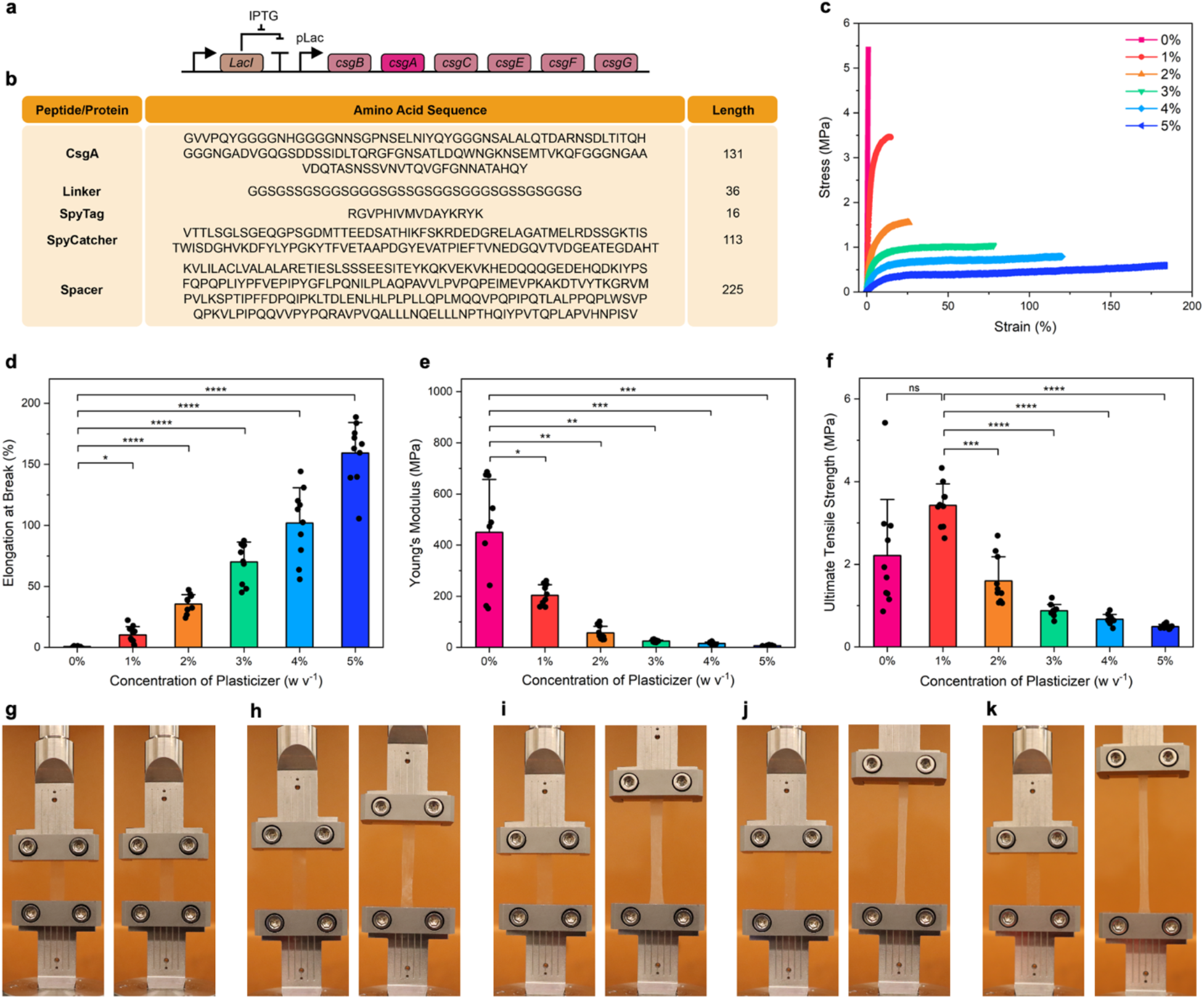
Mechanical Properties of MECHS. a) Genetic design of *E. coli* to produce curli nanofibers. b) Table of amino acid sequences for peptide/protein domains that comprise MECHS variants. c) Representative stress strain curves of MECHS treated with 0 to 5% plasticizer. d) Elongation at break, e) Young’s modulus and f) Ultimate tensile strength of MECHS treated with 0 to 5% plasticizer. *n* = 10. Data represented as mean ± standard deviation. *p ≤ 0.1, **p ≤ 0.01, ***p ≤ 0.001, ****p ≤ 0.0001. One-way ANOVA followed by Tukey’s multiple comparisons test. g-k) Representative photographs of tensile tests of MECHS films with a lateral dimension of 0.5 cm by 4 cm. g) 1%, h) 2%, i) 3%, j) 4%, and k) 5% of plasticizer. Left image: initial. Right image: before break.

### Genetically Engineered Curli Nanofibers to Tailor the Mechanical Properties

Motivated by the above results, we genetically engineered the curli nanofibers to further modulate the mechanical properties of MECHS. We previously developed Biofilm Integrated Nanofiber Display (BIND), wherein genetic fusions to CsgA are used to modulate material properties of assembled curli nanofibers. ^41^ During extracellular self-assembly, the robust *β*-helical blocks of CsgA fusions, stack on top of each other to form functional curli nanofibers with the desired peptide/protein fusions displayed on their surface. We used the genetic programmability of BIND to increase the stiffness of MECHS through covalent crosslinking. To achieve this, we utilized the third generation of split proteins derived from the adhesion domain, CnaB2 of *Streptococcus pyogenes* (SpyTag/SpyCatcher), wherein a spontaneous reaction between the side chains of lysine and aspartic acid residues results in the formation of an isopeptide bond. ^43^ This amide bond formation was reported to have an extremely high reactivity with >90% completion in 15 min at 10 nM concentration, and that for 10 μM, the half-time was less than 30 s. ^43^ Moreover, the reaction does not require any activating groups and is highly specific even in various complex biological media. SpyTag and SpyCatcher were each genetically grafted to CsgA via a linker to obtain CsgA-SpyTag and CsgA-SpyCatcher (**Figure 3a**). ^43^ These two CsgA constructs were expressed from separate plasmids in a co-culture and the resulting curli biomass was used to fabricate MECHS films (denoted as CL1, **Figure 1**). The tensile tests of CL1 showed that their Young’s modulus (51.6 ± 18.4 MPa) and ultimate tensile strengths (1.6 ± 0.4 MPa) were twice that of CsgA only (i.e., not crosslinked) based MECHS films, (**Figure 3c,d,f, S7a** and **S8a**). However, the elongation at break of CL1 was found to reduce to 29.8 ± 8.6% (**Figure 3e**). We also tried analogous experiments with a large spacer (disordered protein domain of 225 amino acids) in between CsgA and the SpyTag/SpyCatcher domains (**Figure 3b** and **Figure 1**). ^44^ MECHS films with this composition (i.e., CL2) were also found to have a Young’s modulus (46.6 ± 27.9 MPa), ultimate tensile strength (1.4 ± 0.7 MPa) and elongation at break (21.9 ± 6%), in the same range as that of CL1 (**Figure 3c-e, S7b** and **S8a,b**). These results clearly indicate that the covalent crosslinking of curli nanofibers in CL1 and CL2 resist the deformation of MECHS films leading to increased Young’s modulus and ultimate tensile strength. However, this was achieved at the expense of elongation at break for CL1 and CL2 films. Although the covalent crosslinks enhance the stiffness of CL1 and CL2, we speculate that the softer biomass in the interstices between curli aggregates provide alternate pathways for crack propagation. Moreover, the slight decrease in the Young’s modulus and the ultimate tensile strength of CL2 in comparison to CL1 might be attributed to the effect of the disordered spacer domain. We reason that an even bigger spacer domain might lead to significant reductions to stiffness and enhanced extensibility.

**Figure 3:**
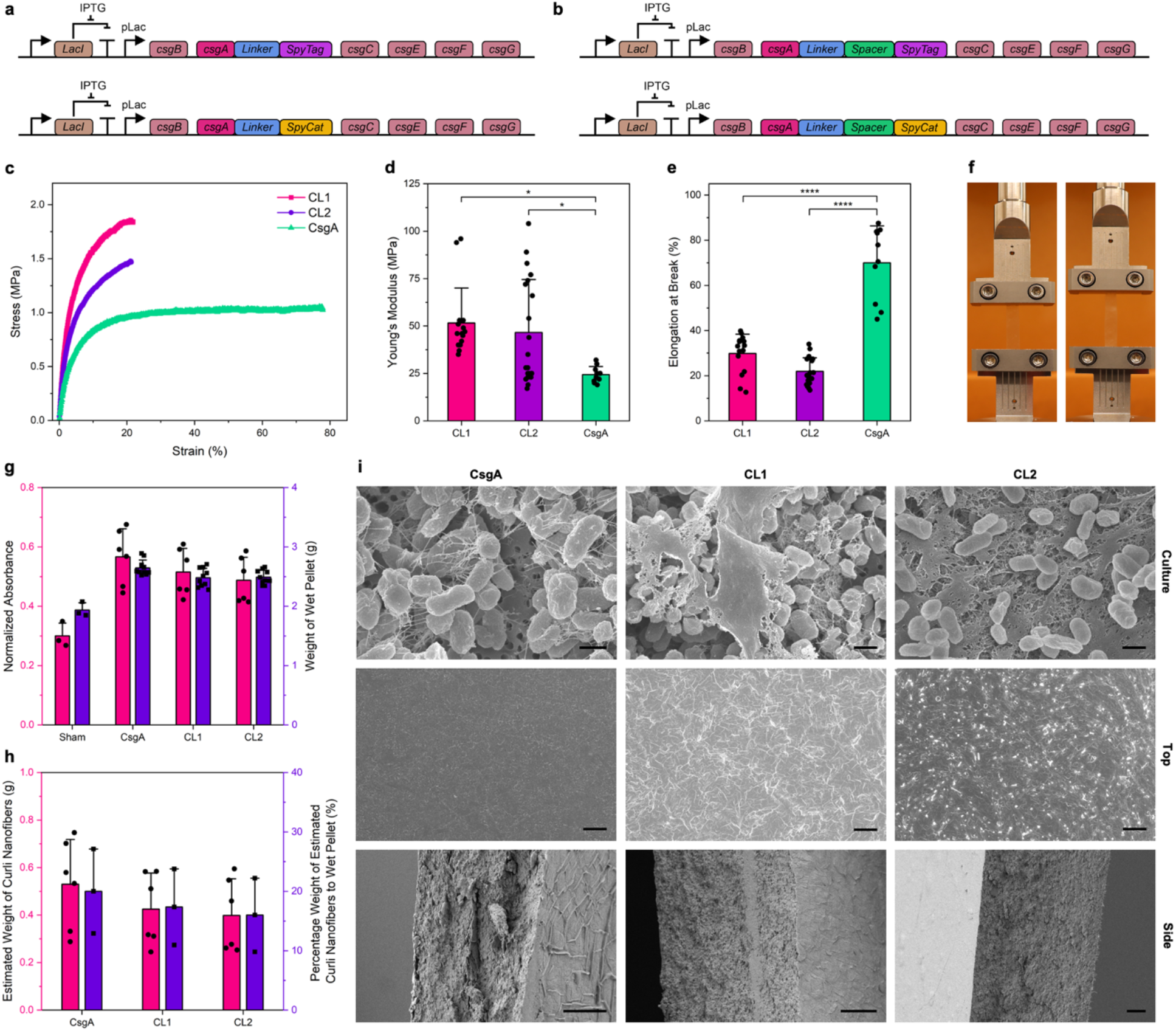
Tailoring the Mechanical Properties of MECHS through genetic engineering. Genetic design of *E. coli* to produce the functional curli nanofibers to covalently crosslink a) CL1: SpyTag and SpyCatcher (SpyCat) domains fused to CsgA, b) CL2: SpyTag and SpyCat domains fused to CsgA via the Spacer. c) Representative stress strain curves of MECHS films consisting of CsgA, CL1 and CL2 with 3% plasticizer. d) Young’s modulus, and e) Elongation at break for CsgA, CL1 and CL2 with 3% plasticizer. *n* = 10. Data represented as mean ± standard deviation. *p ≤ 0.1, ****p ≤ 0.0001. One-way ANOVA followed by Tukey’s multiple comparisons test. f) Representative photographs showing tensile test of CL1 film with the lateral dimension of 0.5 cm by 4 cm. Left image: initial. Right image: before break. g) Plot of normalized Congo Red absorbance and the weights of wet cell pellets. h) Plot of estimated weight of curli nanofibers and the weight percentage of estimated curli nanofibers to the wet pellet. *n* = 3. Data represented as mean ± standard deviation. i) Field Emission Scanning Electron Microscopy (FESEM) images of CsgA, CL1 and CL2. Top row: cell cultures. Scale bar 1 μm. Middle row: Top view of MECHS. Scale bar 10 μm. Bottom row: Side view of MECHS. Scale bar 10 μm.

### Composition and Morphological Analysis

Given the highly heterogeneous nature of the whole biomass that forms MECHS, we wanted to perform a detailed compositional analysis to understand the effects of various components therein. We focused on determining the amounts of curli biomass, gelator, and plasticizer in the final product, which may not be obvious from the fabrication protocol of MECHS. For example, treatment of the wet biomass with 1-5% gelator and/or plasticizer does not mean that the final MECHS film contains 1-5% gelator and/or plasticizer by mass, since only a portion of the original SDS and glycerol will associate with the cell pellet and the rest will be discarded with the supernatant, prior to film casting.

We first focused on estimating the amount of curli nanofibers present in the films on a per-weight basis using a standard Congo Red pull-down assay for curli quantification (**Figure 3g** and **S9a**). These relative amounts of curli were converted to absolute mass estimates with a calibration curve generated using purified curli nanofibers. We estimated that 500 ml cultures of CsgA, CL1, and CL2 produced 530 ± 188 mg, 431 ± 159 mg, and 399 ± 154 mg of curli nanofibers, respectively (**Figure 3h** and **S9b**). The wet weights of whole cell pellets obtained from 500 ml cultures of CsgA, CL1 and CL2 were found to be 2647 ± 130 mg, 2483 ± 157 mg, 2490 ± 118 mg, respectively (**Figure 3g**). Thus, we could estimate the percent of wet weight contributed by curli nanofibers for each construct (**Figure 3h**). Notably, it is possible that the fused SpyTag/SpyCatcher domains may interfere with Congo Red binding, leading to an underestimation of curli nanofibers yields. On the other hand, 500 ml cultures of PQN4 with a sham plasmid (no curli operon) were found to have a wet cell pellet weight of 1936 ± 123 mg (**Figure 3g**). It is interesting to note that the differences in wet pellet mass between curli-producing and sham plasmids roughly corresponds to the mass of curli nanofibers in each culture, calculated from the calibrated Congo Red binding assay (**Figure 3g,h**).

We then set out for an extensive weight analysis to better understand the composition and the effect of various steps involved in the fabrication of MECHS. First, we determined that the ambient drying of the wet pellet of curli biofilm (without the treatment of gelator and plasticizer) results in a dry pellet with a weight percentage (dry to wet pellet) of 20.3 ± 1.8% (**Figure S10a,b**). The dry weight of MECHS films obtained after treatment of 1-5% of gelator was found to be about 100 mg, while the dry weight of the supernatant (collected from all the SDS treatment and water washings of cell pellet) was found to increase linearly (**Figure S11a,b**). It is to be noted that the experimentally obtained sum of weights of MECHS and the corresponding dry supernatant were consistent with their theoretically calculated weights (**Figure S12a-e**). Further, we estimated that the weights of the gelator-treated MECHS films were nearly half of the estimated dry weight (20.3% of wet pellet weight) of curli biomass (**Figure S11c**). Similarly, the weights of MECHS films obtained from 1% and 2% gelator were nearly 45% and 30%, respectively, of the estimated total weight of all precursors, whereas that for 3-5% gelator were about 25% (**Figure S11d**). These results also suggests that the convoluted MECHS films obtained from 1% and 2% gelator upon drying could be attributed to incorporation of more cellular biomass into the films, while the 3-5% gelator might extract more cellular components like lipids into the supernatant (**Figure S2b,c**). Moreover, it is to be noted that unlike 1% and 2% of gelator concentrations, the 3-5% of gelator leads to better gelatinous curli biomass (**Figure S1**). As the percentage weight of MECHS with respect to the dry weight of curli biofilm remains at around 45%, it suggests that the higher gelator (3-5%) content might not lead to additional loss of biomass into supernatant (**Figure S11c**). This latter inference is also supported by the fact that weight of dried supernatant increases in steps of ∼50 mg, which is consistent with the expected increase in the theoretical weights of gelator (*e*.*g*., 5 ml of 1% accounts for 50 mg) (**Figure S11b**).

As noted above, 3-5% gelator-treated MECHS comprises of nearly 45% dry weight of the whole cell pellet, then we reasoned that by determining the amount of SDS, we could estimate the total (cellular and curli) biomass in the MECHS (**Figure S11c**). By using Energy Dispersive X-ray Analysis (EDAX) we found out that for 3% gelator-treated MECHS, the weight percentage of Sodium and Sulfur elements were 2.2 ± 0.2% and 4.5 ± 0.3%, respectively, whereas the same elements for the curli biofilm cell pellet were 0.6 ± 0.1% and 1.2 ± 0.5%, respectively (**Figure S13**). Using this data, we estimate that for 3% gelator-treated films, roughly 5% of MECHS weights could comprise of SDS (**Figure S11a,c**). Therefore, we can estimate that about 40% of the total cellular and curli biomass might get utilized to form the gelator-treated MECHS.

On the other hand, based on the weights of plasticizer-treated MECHS films and their corresponding dry supernatant weights, we could estimate that 15-20 % of the total plasticizer utilized might get incorporated into MECHS, assuming that no additional biomass was lost to the supernatant during this phase of fabrication (**Figure S11, S14** and **S15**). In addition, the weights of MECHS films of CsgA, CL1 and CL2 and their dried supernatants were in the same range, which further validates that the covalent crosslinking in CL1 and CL2, leads to increased stiffness and not due to any variations in the plasticizer amounts (**Figure S16** and **S17**).

Field Emission Scanning Electron Microscopy (FESEM) images from cultures of CL1, and CL2 showed aggregated mats of material, presumably due to nanofiber bundling promoted by the SpyTag/SpyCatcher covalent crosslinking. Images obtained from CsgA cultures did not show such aggregation (**Figure 3i**). FESEM images of MECHS (top and side view) further showed that the curli biomass is densely packed to form continuous films (**Figure 3i, S18** and **S19**).

### Compostability, Scalability and Mechanical Landscape

To test the relative compostability of MECHS films compared to other conventional plastics and bioplastics, we buried samples of each in a commercially available compost called fishnure, derived from fish manure. Experiments were performed in a mini greenhouse setup with samples of uniform size and shape (**Figure S20-21**). Under these conditions, MECHS films biodegraded completely in 15 days, while all the other samples did not (**Figure 4a,c** and **S21-23**). Toilet paper and kimwipes biodegraded to 70% and 40%, respectively (**Figure 4a,c** and **S21**). The bioplastics cellulose acetate (CA) and poly-L-lactic acid (PLLA) were biodegraded by 13% and 1% respectively, whereas the petrochemical plastics polyethylene terephthalate (PET) and low density polyethylene (LDPE) did not show any biodegradation (**Figure S22**). On the other hand, two different commercial polyvinyl alcohol (PVA) formulations, PVA-Mc and PVA-Sp, lost 17% weight and completely disappeared in 5 days, respectively (**Figure S23**).

**Figure 4:**
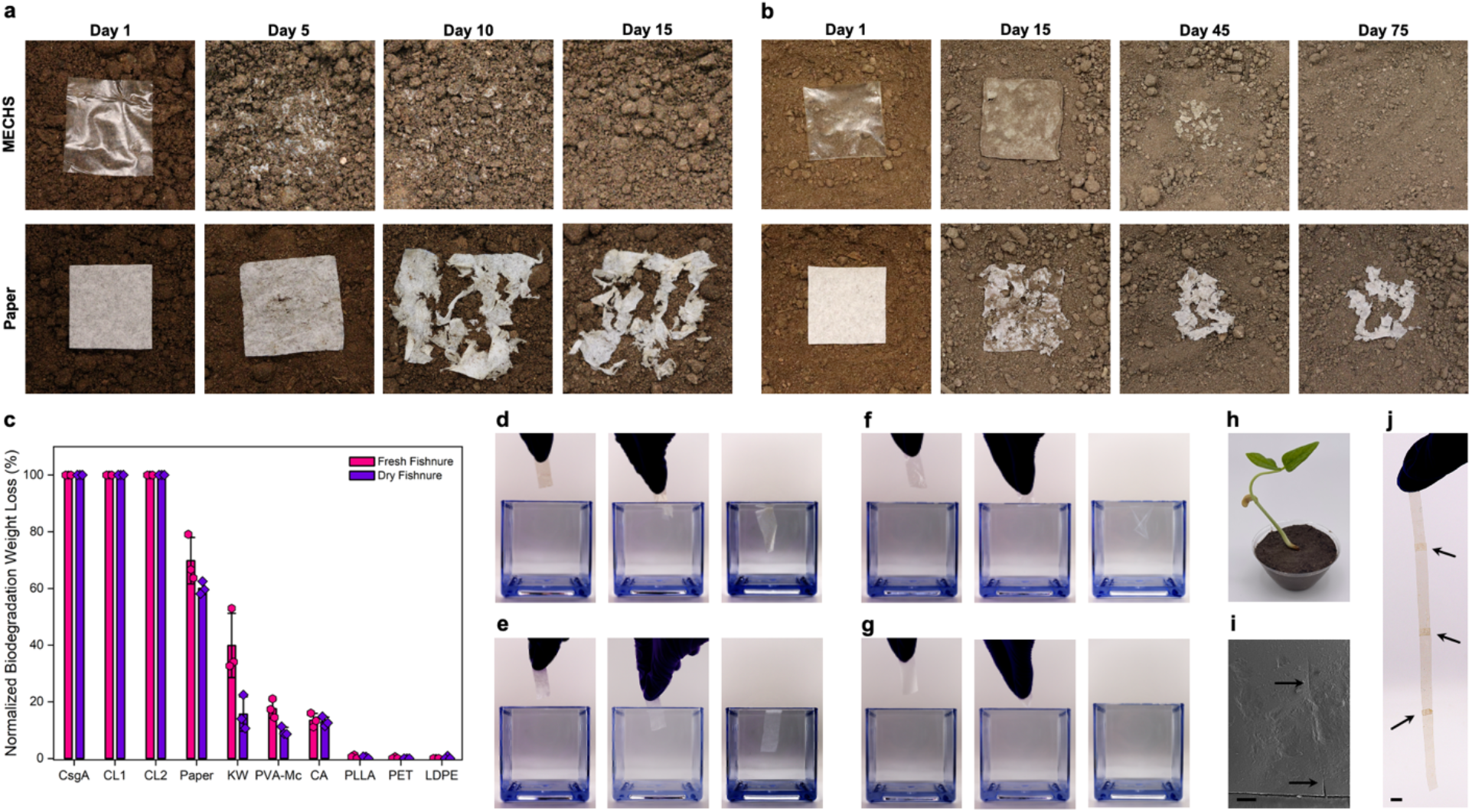
Compostability of MECHS. Representative photographs showing the biodegradation of MECHS and toilet paper in a) a fresh fishnure b) a dry fishnure. The lateral dimensions of the MECHS film and the toilet paper were 5 cm by 5 cm. c) Plot shows the normalized biodegradation weight loss of MECHS (CsgA, CL1 and CL2), toilet paper, kimwipe (KW), polyvinyl alcohol - Mckesson (PVA-Mc), cellulose acetate (CA), poly-L-lactic acid (PLLA), polyethylene terephthalate (PET) and low density polyethylene (LDPE). *n* = 3. Data represented as mean ± standard deviation. Photographs show the dissolution of d) MECHS e) toilet paper, f) PVA-Mc and g) polyvinyl alcohol - Superpunch (PVA-Sp). d-g) Lateral dimension of the films were 1 cm by 5 cm. h) Photograph of a black bean seedling grown in the soil mixed with fishnure (comprising biodegraded MECHS) in 9:1 ratio. i) FESEM image of MECHS film healed by placing microliters of water at the site of abrasion (black arrows). Scale bar 200 μm. j) Photograph shows the MECHS films welded (black arrows) by using water. Scale bar 0.5 cm.

Some of the mass loss in the experiments above may have been attributable to dissolution in the moist fresh fishnure, rather than biodegradation, especially for MECHS and PVA. Therefore, we performed additional compostability tests in fishnure that was dried (i.e., placed in the greenhouse for 50 days). Under these conditions, MECHS films were able to biodegrade completely in 75 days (**Figure 4b,c** and **S24**). The toilet paper, kimwipe and CA were found to degrade by about 60%, 16% and 13%, respectively, whereas PLLA, PET and LDPE did not show any biodegradation in dry fishnure (**Figure 4b,c, S24** and **S25**). However, PVA-Mc had nearly 10% weight loss, whereas PVA-Sp was found to be intact even after 75 days in dry fishnure. We could not determine the weight loss of PVA-Sp as the film was firmly sticking to the fishnure granules. These experiments show that the MECHS films are completely compostable and that their biodegradation compares favorably to many plastics, bioplastics and even toilet paper.

For potential use of MECHS as flushable packaging materials, we tested its ability to dissolve in water (**Figure 4d,e**). The MECHS films did not dissolve completely, likely due to the network of hydrophobic curli nanofibers. We speculate that the more water-soluble components like glycerol, SDS and the other cellular biomass leach into water more readily. PVA-Sp dissolves completely in water whereas PVA-Mc dissolves only partially, leaving behind water-insoluble strips possibly compromising its biodegradation (**Figure 4f,g**). Furthermore, except for MECHS, all the other plastics compared here are composed only of carbon, hydrogen, and oxygen. Therefore, their biodegradation is often considered, in terms of breakdown completely, to lead to carbon dioxide. However, MECHS is largely composed of protein, making it the only plastic amongst those compared here with any significant nitrogen content. Therefore, it may be reasonable to consider its potential as a biofertilizer to support plant growth (**Figure 4h**). Further, MECHS films could also be healed and welded by using microliters of water at the site of abrasion or attachment and subsequently ambient dried (**Figure 4i,j**).

The fabrication method presented in this work yielded 500 to 1000 mg of MECHS films per liter of culture, which is nearly 10 times higher than the 50 to 100 mg obtained from our previously reported “AquaPlastic” protocol. ^9^ We achieved these yields even with a standard shake-flask format that is routinely used in laboratory settings for recombinant protein production. Therefore, tens of liters of bacterial culture could be used to fabricate large MECHS prototypes, such as thin films tens of centimeters in one dimension (**Figure 5a,b** and **S26**). We also created a detergent pod as an example of flushable and biodegradable primary package (**Figure 5c**).

**Figure 5:**
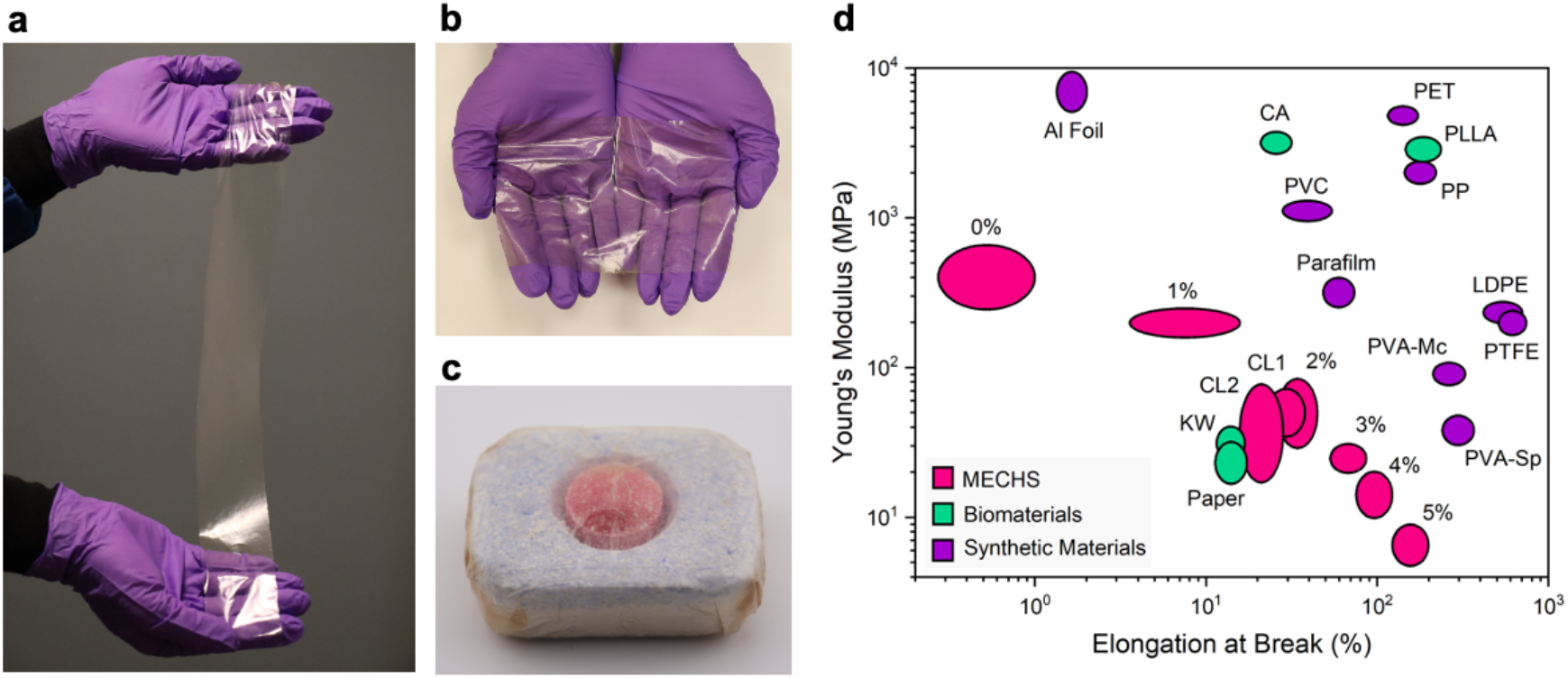
Prototypes and Mechanical Landscape of MECHS. a) Photograph shows a refined prototype of MECHS film with a lateral dimension of 5 cm by 50 cm. b) Photograph shows the optical transparency of MECHS film with a lateral dimension of 10 cm by 15 cm. c) Photograph of a detergent pod (lateral dimension of 4 cm by 3 cm) wrapped with a MECHS film. d) Ashby plot shows the Young’s modulus and elongation at break for MECHS and various synthetic materials and biomaterials. Low density polyethylene (LDPE), Polytetrafluoroethylene (PTFE), Poly-L-lactic acid (PLLA), Polyethylene terephthalate (PET), Cellulose acetate (CA), Polypropylene (PP), Polyvinyl chloride (PVC), Polyvinyl alcohol - Superpunch (PVA-Sp), Polyvinyl alcohol - Mckesson (PVA-Mc), Aluminum foil (Al Foil), Parafilm, Kimwipes (KW) and Toilet paper.

To better visualize and compare the mechanical properties of MECHS, we present an Ashby plot of Young’s modulus and elongation at break for various plastics, bioplastics, biomaterials, and synthetic materials (**Figure 5d, S27-35**). It is thus evident that the Young’s modulus of MECHS is in the same range of LDPE, PTFE (polytetrafluoroethylene), PVA, and paper, while its elongation at break matches that of CA, PVC (polyvinyl chloride), PP (polypropylene), PET, PLLA and parafilm.

## Discussion and Conclusion

We previously reported the curli nanofiber-based bioplastic fabrication protocol (i.e., AquaPlastic), which involved the filtration of bacterial culture to concentrate curli nanofibers and form gels. ^9^ Using that protocol, concerns about clogging necessitated the use of filters with 10 μm pores, leading to the loss of significant amounts of curli nanofibers. The MECHS fabrication protocol described in this paper increased the yield of bioplastic by a factor of ten by utilizing not only all the curli nanofibers in the pelletized biomass, but also the other water insoluble cellular biomass. We also found that the SDS gelator could be supplemented with a plasticizer like glycerol to obtain flexible films of MECHS, as compared to the significantly more brittle AquaPlastic. Glycerol being a byproduct of the biodiesel industry offers several advantages *viz*., nontoxic, low-cost, and renewable. ^45^ Unlike the conventional petrochemical plastics and other bioplastics that are processed by thermal molding, MECHS was molded into films by ambient drying of gelatinous biomass, which we have termed it as aquamolding. The healing and welding of MECHS films by using tiny droplets of water are termed as aquahealing and aquawelding, respectively.

The tunability of MECHS, with its range of mechanical properties (e.g., elongation at break 1-160%; Youngs’ modulus 6-450 MPa) and transparency, provides a promising platform to access biodegradable alternatives to synthetic materials like petrochemical plastics. We were also able to use our streamlined protocol to achieve high yields of 0.5-1 g L^-1^ and generate large, refined prototypes. Another notable feature of this work is that 40% of the total cellular biomass gets incorporated into the plastic/paper-like MECHS, which could also be instrumental in attracting further research to utilize cellular biomass for the development of various sustainable functional materials.

Plastics are one of the most abundant human-made materials, with over 8.3 billion tons produced cumulatively, 79% of which are estimated to have accumulated in landfills and oceans. ^46^ In addition, the contamination of microplastics in almost all parts of the globe further enhances their threat to our health and the environment. ^47,48^ Biodegradable bioplastics account for less than 1% of the global plastic market and their limited properties warrants the development of new alternatives. ^35^ Given that the typical lifetime of a packaging material is 1-2 years, and the packaging industry accounts for nearly one third of the plastic market, there exists a tremendous scope and opportunity for biodegradable packaging, though success will likely need to be achieved through the commercialization of drop-in replacements for existing materials. Notably, water soluble polymers like PVA (commonly found in detergent pods) have limited biodegradation under diverse settings of land and water. ^49^ In many cases, dissolvable polymers like PVA are blended with petrochemical plastics to enhance certain material properties but this limits their water dispersibility and biodegradability (as observed in our biodegradation tests with the commercially available PVA-Mc). ^50^

Although we were able to develop refined prototypes of MECHS thin films, additional work will be needed to improve the mechanical properties (e.g., ultimate tensile strength, tear strength) and resistance to water. Furthermore, the circular materials economy loop will have to be closed by employing a feedstock for bacterial culture derived closely from CO_2_ fixation, such as cellulose hydrolysate obtained from agricultural waste. There are also several opportunities to utilize the synthetic biology tools to tailor the material properties of curli nanofibers, which needs to be explored. The concept of using biodegraded MECHS as a biofertilizer for plant growth warrants further investigation. All in all, in this work we have demonstrated that the manufacturing capabilities of living cells can be employed to produce the mechanically tunable, scalable, and compostable ELMs as a potential alternative to synthetic materials like plastics. Finally, we believe that innovative approaches involving synthetic biology and materials engineering could lead to greater advancements in creating energy efficient and sustainable solutions to a greener ecosystem.

## Acknowledgements

Work was performed in part at the Center for Nanoscale Systems at Harvard University and George J. Kostas Nanoscale Technology and Manufacturing Research Center, Northeastern University. Work in the N.S.J. laboratory is supported by the National Science Foundation, USA (DMR 2004875) and Novo Nordisk Foundation Challenge Programme 2022 - Energy Materials with Biological Applications (NNF22OC0071130). We thank Arjun Rajesh and Bismay Hota for their assistance with capturing photographs of MECHS biodegradation and prototypes. Parts of the schematics were adapted from BioRender.com.

## Author contributions

A.M.-B. conceived the project and performed all the experiments and analysis. A.M.-B. and A.M.D.-T. cloned all the curli variants. A.M.-B., and N.S.J. wrote and edited the manuscript. All authors discussed and commented on the manuscript.

## Competing interests

A.M.-B., A.M.D.-T., and N.S.J. are inventors on a U.S. Provisional Patent Application (63/604,497) submitted by Northeastern University.

## Data availability

All relevant data supporting the findings of this study and the plasmids and strains used are available within the Article and its Supplementary Information or from the corresponding authors upon request. Source data are provided with this paper.

## Notes

### Competing Interest Statement

Avinash Manjula-Basavanna, Anna M. Duraj-Thatte and Neel S. Joshi are inventors on a U.S. Provisional Patent Application (63/604,497) submitted by Northeastern University.

