## Supplementary Information for "Mechanically Tunable, Compostable, Healable and Scalable Engineered Living Materials"

### Materials and Methods

#### Materials

Low density polyethylene LDPE (ET31-FM-000151, 50 μm thick), Polytetrafluoroethylene PTFE (FP30-FM-000250, 50 μm thick), Poly-L-lactic acid PLLA (ME33-FM-000150, 50 μm thick), Polyethylene terephthalate PET (ES30-FM-000150, 50 μm thick), Cellulose acetate CA (AC31-FM-000151, 50 μm thick) and Polypropylene PP (PP30-FM-000250, 50 μm thick) were obtained from Goodfellow Corporation. Polyvinyl chloride PVC (S-16280, 15 μm thick) was obtained from Uline. Polyvinyl alcohol - Superpunch PVA - Sp (ASIN: B01M11T6U5), Polyvinyl alcohol - Mckesson PVA - Mc (ASIN: B01ETFMUH2) and Silicone mats (ASIN: B09SPB72TT) were obtained from Amazon. Aluminum foil, Parafilm, Toilet paper and Kimwipes were obtained from Reynolds Consumer Products, Bemis Company Inc., Signature Select and Kimberly-Clark Corporation, respectively. Glycerol (G9012) and Sodium dodecyl sulfate SDS (S0295) were obtained from Sigma-Aldrich and Teknova, respectively.

**Plasmids to produce MECHS:** pET21d plasmid was cloned with the curli operon genes *csgA*, *csgB, csgC*, *csgE*, *csgF* and *csgG* that encodes the proteins necessary for biosynthesis of curli nanofibers and it is labelled as pET21d-CsgA. The genes encoding the SpyTag peptide and SpyCatcher protein derived from an earlier report[1] were fused to the C-terminus of CsgA with an intervening 36 amino acid flexible linker to obtain the plasmids pET21d-CsgA-SpyTag and pET21d-CsgA-SpyCatcher, respectively. The gene encoding the Spacer, an intrinsically disordered protein[2], was inserted between the linker and the SpyTag or the SpyCatcher to obtain pET21d-CsgA-Spacer-SpyTag and pET21d-CsgA-Spacer-SpyCatcher, respectively. The genes were synthesized (Integrated DNA Technologies) and cloned into pET21d vector using isothermal Gibson assembly (New England Biolabs).

**Cell Strain to produce MECHS:** The plasmids pET21d-CsgA, pET21d-CsgA-SpyTag, pET21d-CsgA-SpyCatcher, pET21d-CsgA-Spacer-SpyTag and pET21d-CsgA-Spacer-SpyCatcher were separately transformed into PQN4, an *E. coli* cell strain derived from LSR10 (MC4100, Δ*csgA*, λ(DE3), $Cam^R$) with the deletion of curli operon (ΔcsgBACEFG) to produce the corresponding MECHS.[3]

**Cell Culture to produce MECHS (CsgA):** pET21d-CsgA plasmid was transformed into PQN4 and streaked onto lysogeny broth (LB) agar plate containing 100 μg $ml^{-1}$ carbenicillin and 0.5% glucose (m $v^{-1}$) for catabolite repression of T7RNAP and incubated overnight at 37 °C. A single colony of PQN4-pET21d-CsgA was picked from the agar plate and cultured at 37 °C in 5 ml LB media, 100 μg $ml^{-1}$ carbenicillin and 2% glucose (m $v^{-1}$). The overnight culture was transferred to a fresh 500 ml LB media containing 100 μg

$ml^{-1}$ carbenicillin and cultured for 48 h in incubator shakers (225 rpm, 37 °C) to express the CsgA curli protein nanofibers.

**Cell Culture to produce the Covalently Crosslinked MECHS (CL1 and CL2):** For Covalently Crosslinked-1 (CL1), the plasmids pET21d-CsgA-SpyTag and pET21d-CsgA-SpyCatcher were separately transformed into PQN4 and streaked onto lysogeny broth (LB) agar plates containing 100 μg $ml^{-1}$ carbenicillin and 0.5% glucose (m $v^{-1}$) for catabolite repression of T7RNAP and incubated overnight at 37 °C. A single colony was picked from the agar plates of PQN4-pET21d-CsgA-SpyTag and PQN4-pET21d-CsgA-SpyCatcher and cultured separately at 37 °C in 5 ml LB media, 100 μg $ml^{-1}$ carbenicillin and 2% glucose (m $v^{-1}$). The overnight cultures of PQN4-pET21d-CsgA-SpyTag and PQN4-pET21d-CsgA-SpyCatcher were transferred to a fresh 500 ml LB media containing 100 μg $ml^{-1}$ carbenicillin and co-cultured for 48 h in incubator shakers (225 rpm, 37 °C) to express and covalently crosslink the engineered curli protein nanofibers. Similarly, the plasmids pET21d-CsgA-Spacer-SpyTag and pET21d-CsgA-Spacer-SpyCatcher were utilized for Covalently Crosslinked-2 (CL2) MECHS.

**Biofabrication of MECHS:** The 48 h cell culture (500 ml) of PQN4-pET21d-CsgA (CsgA) was centrifuged (5000 rpm, 10 min) to pelletize the curli biomass, which was then washed with 250 ml of deionized water by centrifuging (5000 rpm, 10 min) to remove the residual quantities of culture media. 1 g (wet pellet) of curli biofilm biomass was first dispersed in 5 ml of deionized water and subsequently added with 5 ml of 1%, 2%, 3%, 4% or 5% (w $v^{-1}$) of sodium dodecyl sulfate (SDS, serves as a gelator), which was then mixed on a shaker for 2 h at room temperature. The resulting gelatinous biomass was washed with 10 ml of deionized water twice by centrifuging (5000 rpm, 10 min) to remove the soluble biomolecules and the excess SDS. This SDS treated gelatinous biomass was casted and ambient dried on a silicone mold to obtain the MECHS films that were extremely brittle.

To realize the flexible films of MECHS, the 3% SDS treated gelatinous biomass of PQN4-pET21d-CsgA (CsgA) was added with 5 ml of 1%, 2%, 3%, 4%, or 5% (w $v^{-1}$) of glycerol (serves as a plasticizer) and mixed on a shaker for 1 h at room temperature. The glycerol treated and centrifuged (5000 rpm, 10 min) biomass was casted on a silicone mold and ambient dried to obtain the flexible MECHS films.

Similarly, to realize the Covalently Crosslinked (CL1 and CL2) films of MECHS, 5 ml of 3% SDS and 5 ml of 3% glycerol treated curli biomass was utilized. For all constructs, a minimum of ten replicates were reported.

**Field-emission scanning electron microscopy (FESEM) sample preparation and imaging:** 100 μL of cell culture was vacuum filtered on a membrane (0.22 μm pore size, Millipore GTTP02500) and washed with 100 μL of deionized water thrice. The samples were fixed by immersing in 2 ml 1:1 mixture of 2% (w $v^{-1}$) glutaraldehyde and 2% (w $v^{-1}$)

paraformaldehyde at room temperature, overnight. The samples were gently washed with water, and the solvent was gradually exchanged to ethanol (200 proof) with an increasing ethanol 15-minute incubation step gradient [25, 50, 75 and 100% (v $v^{-1}$) ethanol]. The samples were then dried in a critical point dryer, placed onto SEM sample holders using silver adhesive (Electron Microscopy Sciences) and sputtered until they were coated in a 10-20 nm layer of Pt/Pd. Whereas the films of MECHS were directly sputter coated with 10-20 nm layer of Pt/Pd without critical point drying. Images were acquired using a Zeiss Gemini 360 FESEM equipped with a field emission gun operating at 5-10 kV. Representative images from three independent samples were reported.

**Energy Dispersive X-ray Analysis (EDAX):** Oxford Instruments Ultim Max EDS attached to Zeiss Gemini 360 FESEM was utilized to detect the elements as well as determine their composition using factory standards. EDS spectra were recorded on sample's surface with the lateral dimensions of 225 µm by 170 µm. Data from three independent samples were reported.

**Optical Images:** Optical images were acquired using a Canon EOS Rebel SL3 Digital SLR Camera equipped with XIT 58 mm 0.43 Wide Angle Lens and XIT 58 mm 2.2x Telephoto Lens. Representative images from three independent samples were reported.

**Tensile Tests:** Tensile measurements of MECHS, commercially available plastics, bioplastics, and all other materials mentioned in this report were performed using a DHR-3 rheometer (TA Instruments) under ambient laboratory conditions. Films with the lateral dimensions of 4 cm by 0.5 cm under a constant linear deformation of 1 µm $s^{-1}$ were utilized for tensile tests. A minimum of five samples were tested for each type.

**Film Thickness:** The thickness of the films was measured using a contact profilometer, Dektak 3ST equipped with a 2.5 µm stylus having a vertical resolution of 1Å. A minimum of three tests were performed for each sample.

**Large Prototypes:** The MECHS prototype of 50 cm × 5 cm lateral dimension was fabricated from 6 L cultures of PQN4-pET21d-CsgA (obtained by using 3% SDS and 3% glycerol treatment), whereas the 15 cm × 10 cm and the detergent pod prototypes were obtained from that of 4 and 3 L cultures, respectively.

**Healing:** The films of MECHS (PQN4-pET21d-CsgA, obtained by using 3% SDS and 3% glycerol) were cut using scissors and ~10 µL of deionized water was added at the cut site and subsequently dried at ambient laboratory conditions to heal the cut. A minimum of three samples were tested. Similarly, MECHS films of 0.5 cm by 5 cm were welded by using ~10 µL of deionized water and subsequently dried at ambient laboratory conditions.

**Biodegradation:** A commercially available odorless organic humus compost named Fishnure (Amazon, ASIN: B086KXT5TQ), which is made from fish manure was utilized for the biodegradation test. Samples with the lateral dimensions of 5 cm × 5 cm were buried in a tray containing 3.5 kg of Fishnure. The biodegradation experiment was conducted in a mini greenhouse (Amazon, ASIN: B01D7GHEES) setup (exposed to direct/indirect sunlight through the large windows of the laboratory), wherein a temperature of 20 °C and a relative humidity of 80% was maintained. The films of MECHS degraded completely in 15 days in a freshly opened bag of Fishnure. In another biodegradation experiment, a dry (by placing in the mini greenhouse setup for 50 days) Fishnure was utilized and under these conditions, films of MECHS degraded completely in 75 days. A minimum of three samples were tested for each type.

**Congo Red assay**: 1 ml of cell culture (as described above: 48 h, 500 ml at 37 °C) was pelleted by centrifuging (6000 rpm, 10 min) and the resulting cell pellet was incubated with 1 ml of 0.004% (w $v^{-1}$) Congo Red dye for 10 min. The dye treated cell culture was pelletized by centrifuging (6000 rpm, 10 min) and the resulting supernatant (200 $\mu$L) was utilized to measure the absorbance at 480 nm in a plate reader. To estimate the curli nanofibers produced, we utilized 0.004% (w $v^{-1}$) Congo Red dye to prepare a standard curve for various concentrations of purified CsgA. A minimum of three samples were tested for each type.

**Statistics and Reproducibility:** All experiments presented in this Article were repeated at least three times (n ≥ 3) on distinct samples, as clearly specified in the figure legends or the relevant Methods sections. In all cases, data are presented as the mean and standard deviation. OriginPro and GraphPad Prism 8 software were used for plotting and analyzing data. For micrographs and optical images, we present representative images.

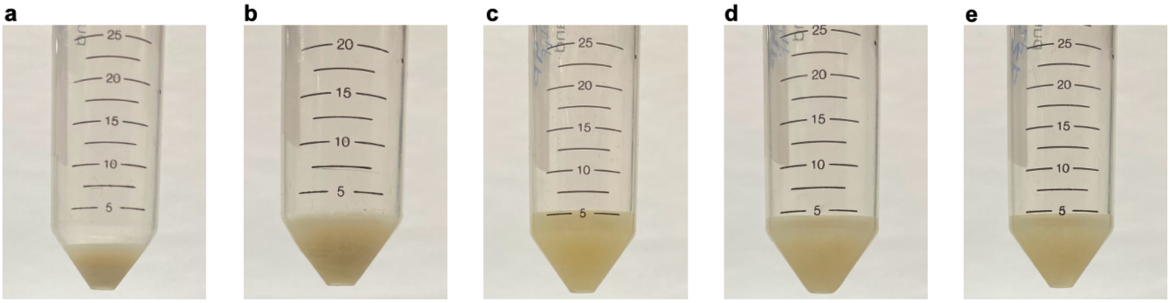


**Figure S1: Gelator treated curli biomass.** Photographs of curli biomass obtained from a) 1%, b) 2%, c) 3%, d) 4% and e) 5% gelator (sodium dodecyl sulfate).

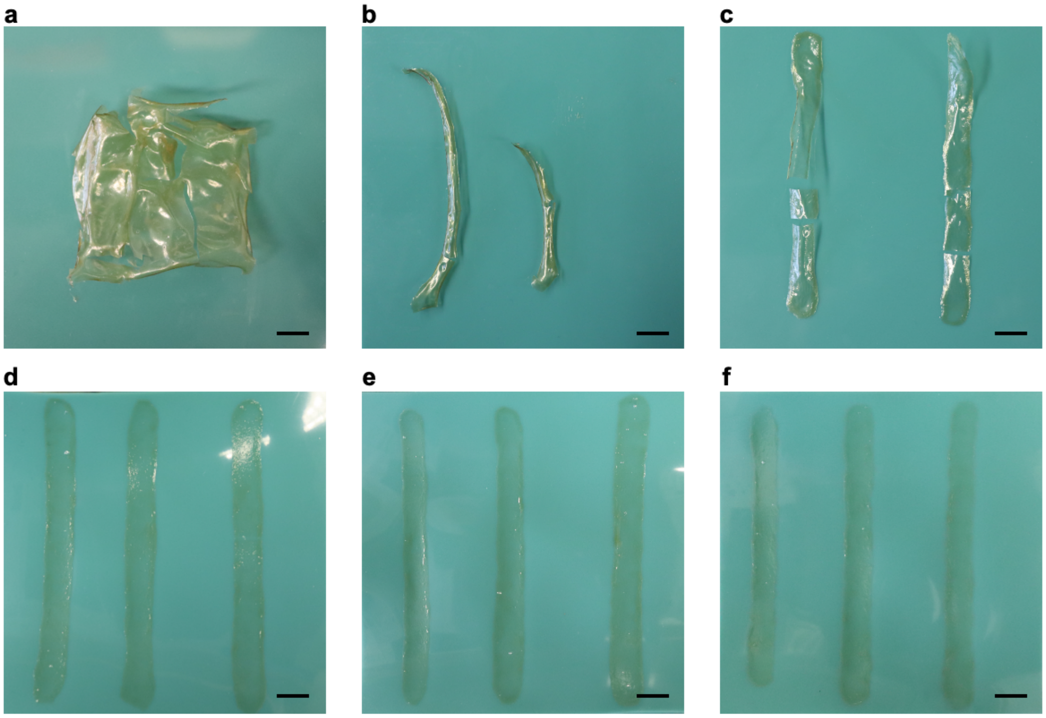


**Figure S2: Gelator treated MECHS.** Photographs of MECHS obtained from a) 0%, b) 1%, c) 2%, d) 3%, e) 4% and f) 5% gelator (sodium dodecyl sulfate). Scale bar 1 cm.

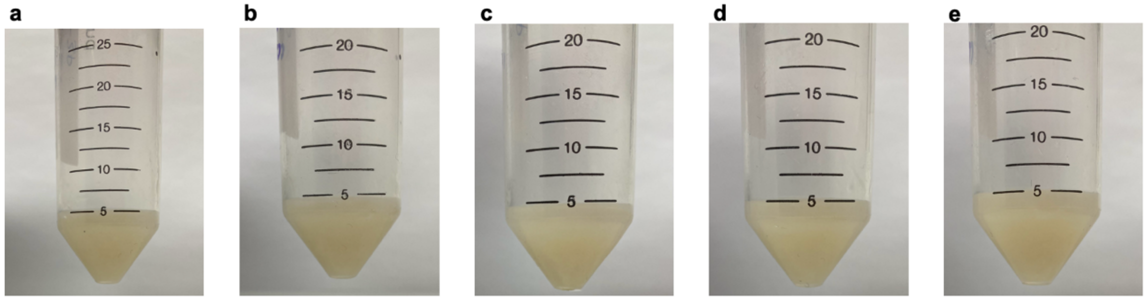


**Figure S3: Plasticizer treated curli biomass.** Photographs of curli biomass obtained from a) 1%, b) 2%, c) 3%, d) 4% and e) 5% plasticizer (glycerol).

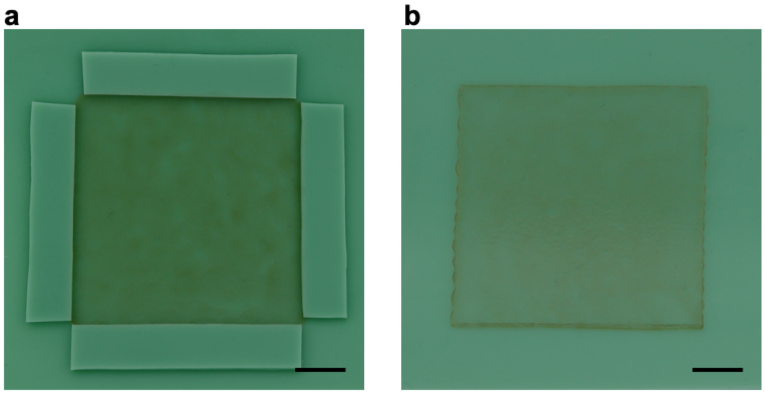

**Figure S4: Plasticizer treated MECHS.** a) and b) Photographs of MECHS film obtained by treating with 3% plasticizer (glycerol) and casting on a silicone mold. Scale bar 1 cm.

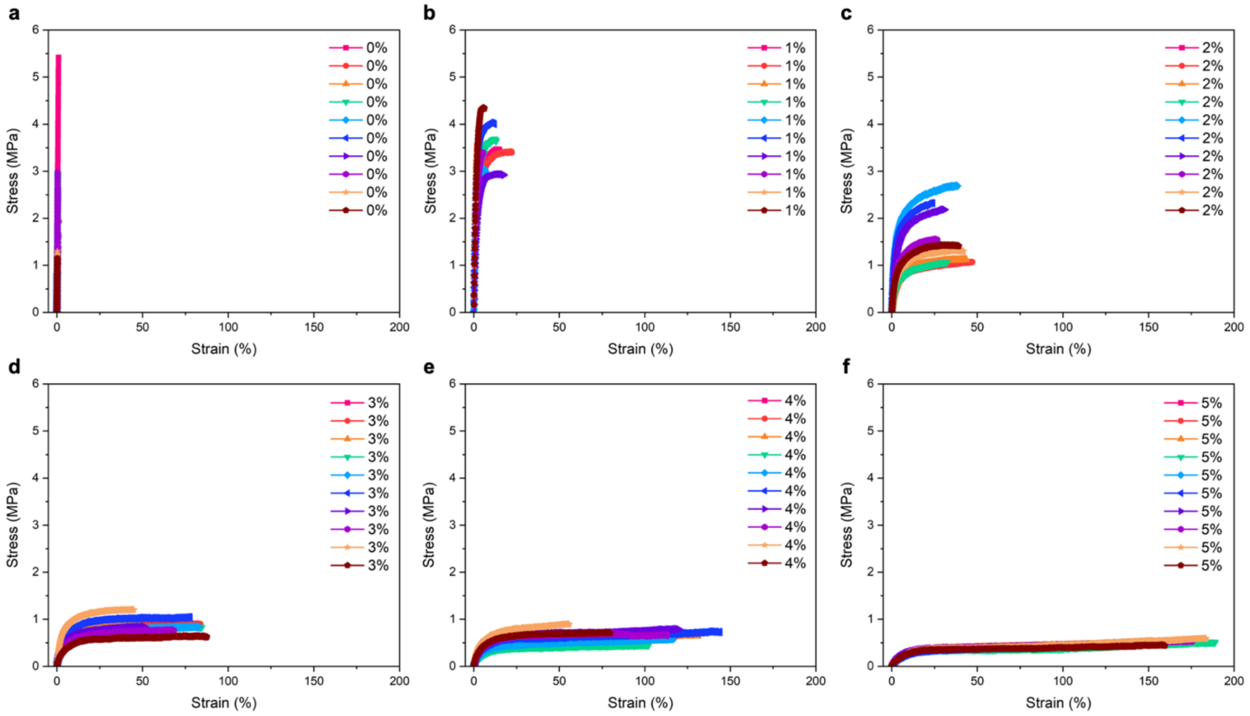


**Figure S5: Tensile tests of MECHS.** Stress strain curves of MECHS films obtained from a) 0%, b) 1%, c) 2%, d) 3%, e) 4% and f) 5% plasticizer (glycerol).

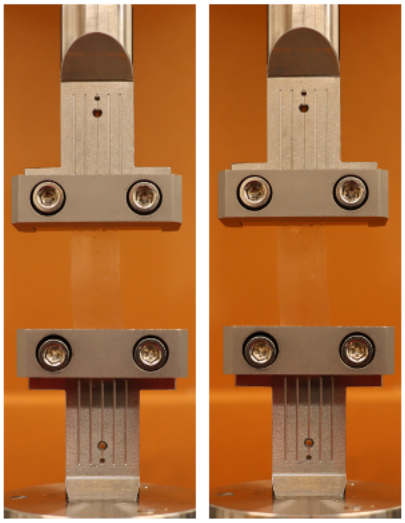

**Figure S6: Tensile tests of MECHS.** Representative photographs show the tensile tests of MECHS film with the lateral dimension of 1 cm by 4 cm obtained from 3% of gelator. Left image: initial. Right image: before break

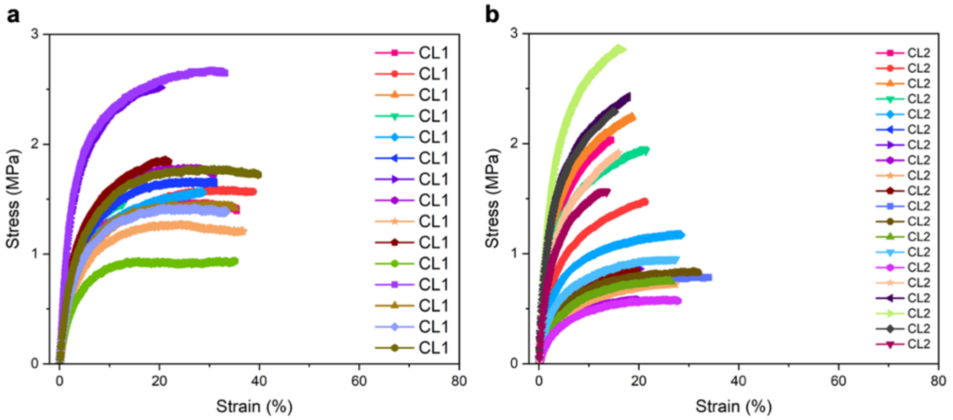


**Figure S7: Tensile tests of MECHS.** Stress strain curves of MECHS obtained from a) CL1 and b) CL2.

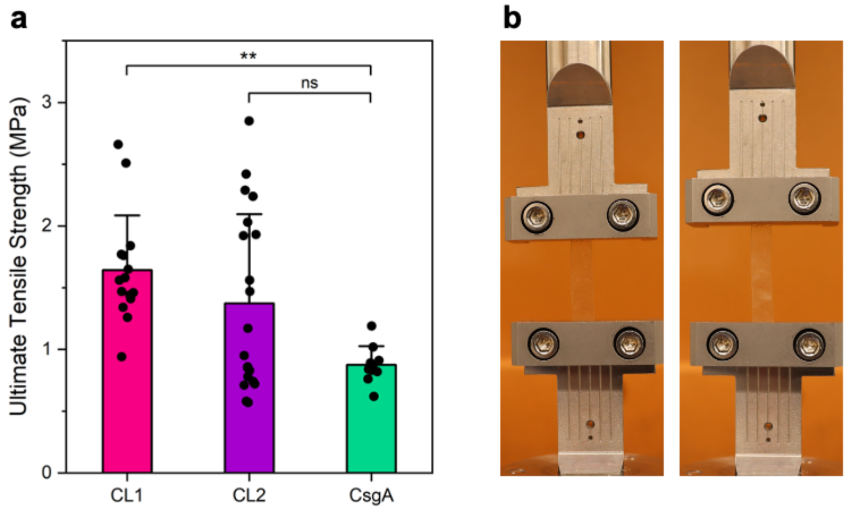


**Figure S8: Tensile tests of MECHS.** a) Ultimate tensile strength of CsgA, CL1 and CL2 with 3% plasticizer. $n \geq 10$. Data represented as mean ± standard deviation. **p ≤ 0.01. One-way ANOVA followed by Tukey's multiple comparisons test. b) Representative photographs show the tensile test of CL2 film with the lateral dimension of 0.5 cm by 4 cm. Left image: initial. Right image: before break.

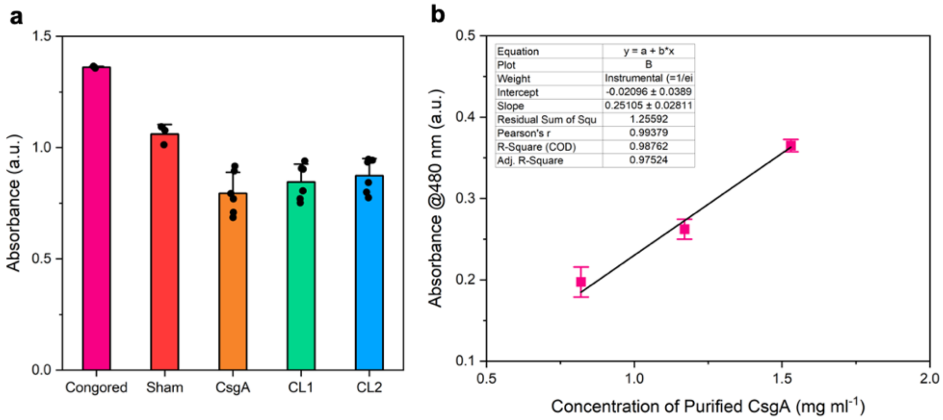


**Figure S9: Congo Red assay.** a) Congo Red absorbance at 480 nm for various samples. b) Congo Red standard curve for purified CsgA. $n \geq 3$. Data represented as mean ± standard deviation.

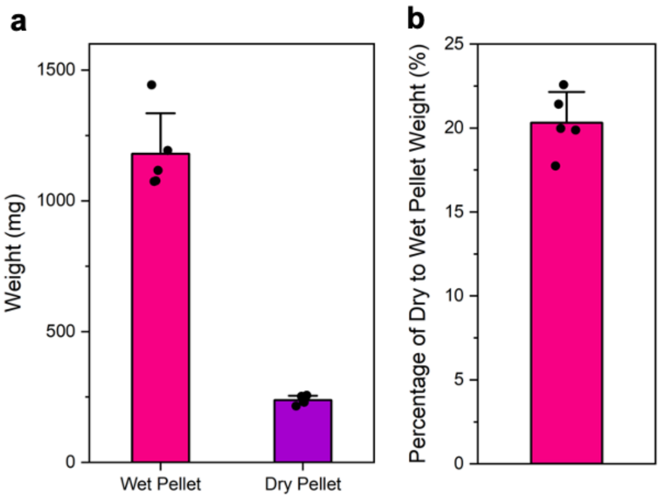


**Figure S10: Weight analysis.** a) Weight of wet and dry pellet of curli biofilm obtained from 500 ml cultures. b) Percentage of dry to wet pellet weight of curli biofilm. *n* = 5. Data represented as mean ± standard deviation.

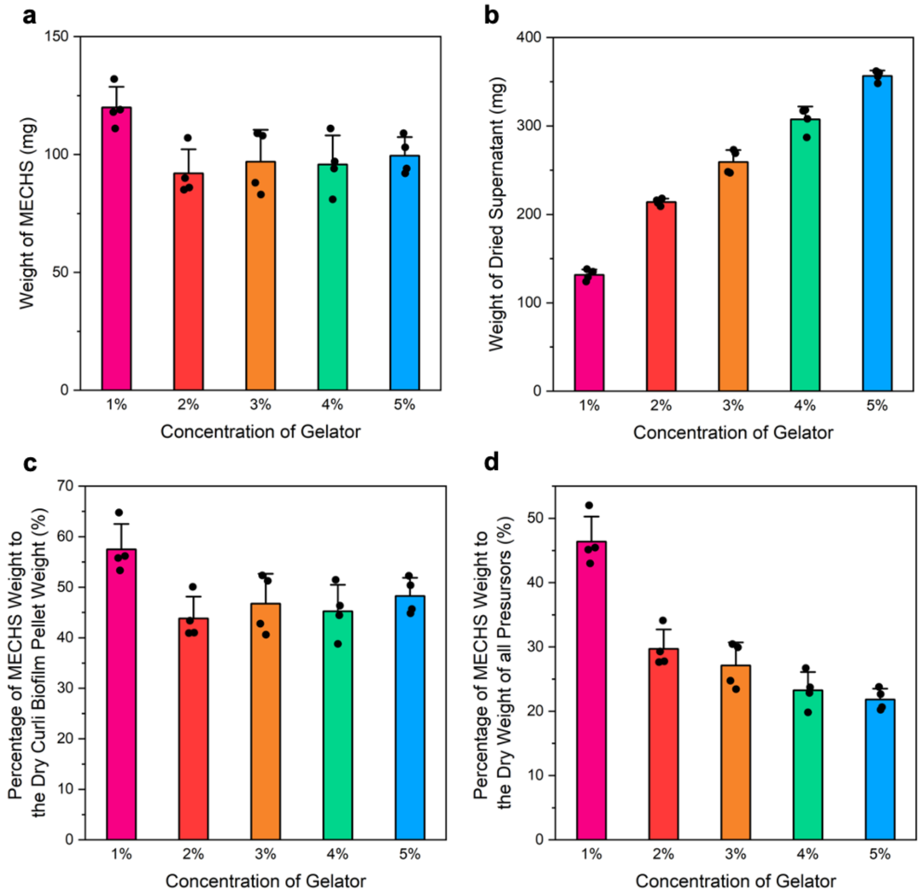


**Figure S11: Weight analysis of gelator treated MECHS.** Weight of a) MECHS films and b) the dried supernatant obtained from 1 to 5% of gelator. Percentage of MECHS film weight to c) dry curli biofilm pellet weight (20.3% of wet pellet) and d) dry weight of all precursors for 1 to 5% of gelator. All precursors correspond to weights of cell pellet and that of 1-5% gelator. *n* = 4. Data represented as mean ± standard deviation.

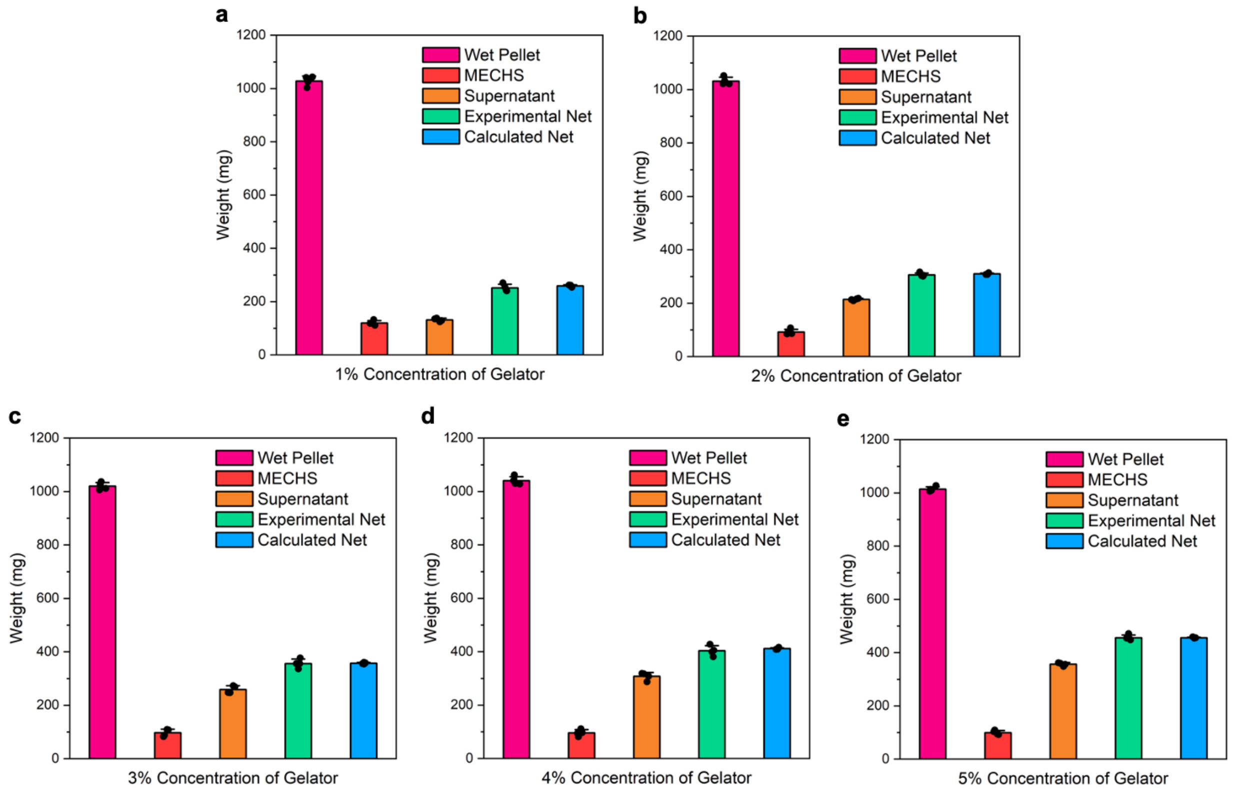


**Figure S12: Weight analysis of gelator treated MECHS.** Weights of wet pellet, MECHS film, dried supernatant, experimental net (MECHS and dried Supernatant) and calculated theoretical net (20.3% of wet pellet) for a) 1%, b) 2%, c) 3%, d) 4% and e) 5% of gelator. $n$ = 4. Data represented as mean ± standard deviation.

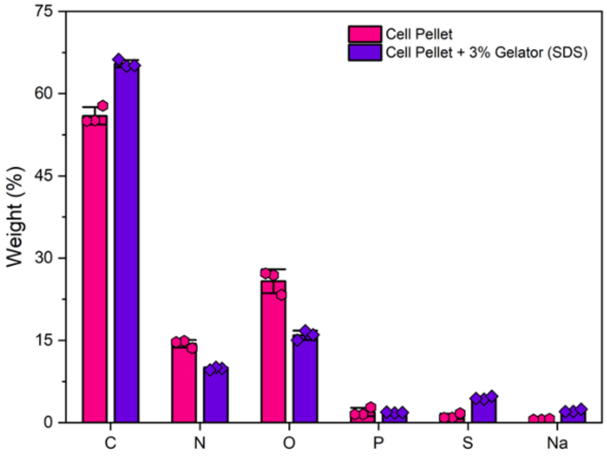


**Figure S13: Compositional analysis of MECHS.** Energy Dispersive X-ray Analysis (EDAX) of *E. coli* curli biofilm cell pellets pretreated with or without 3% gelator (SDS) shows the weight percentage of various elements. $n$ = 3. Data represented as mean ± standard deviation.

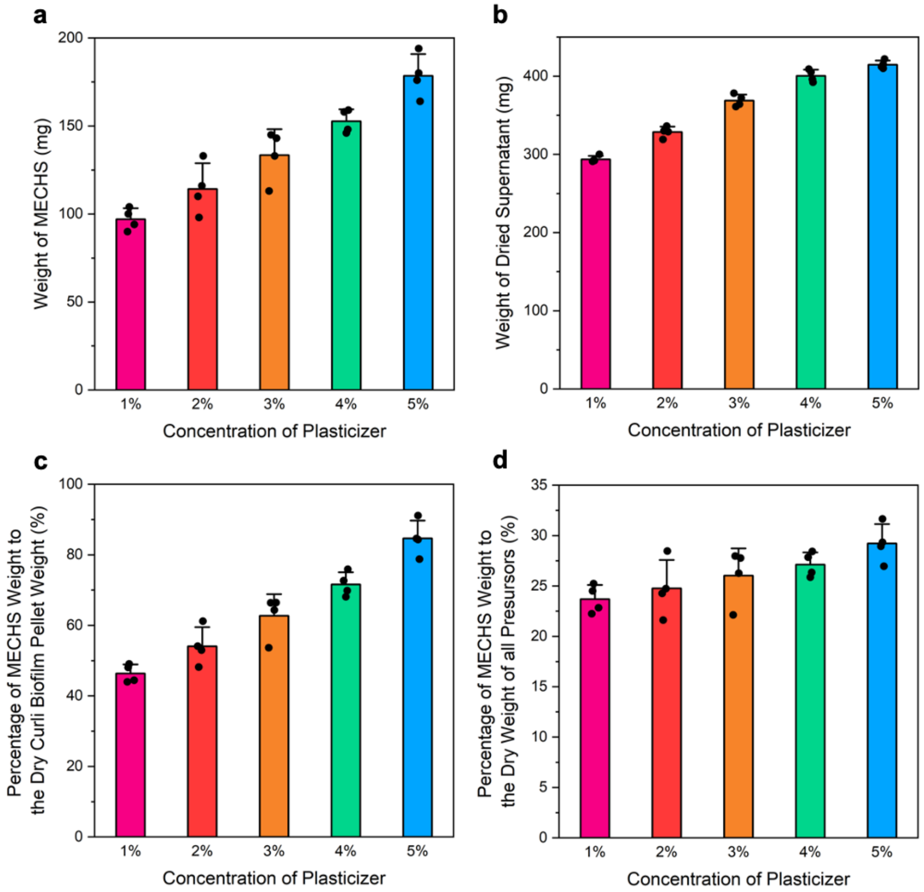


**Figure S14: Weight analysis of plasticizer treated MECHS.** Weight of a) MECHS films and b) the dried supernatant obtained from 1 to 5% of plasticizer. Percentage of MECHS film weight to c) dry curli biofilm pellet weight (20.3% of wet pellet) and d) dry weight of all precursors for 1 to 5% of plasticizer. All precursors correspond to weights of cell pellet and that of 3% gelator and 1-5% plasticizer. *n* = 4. Data represented as mean ± standard deviation.

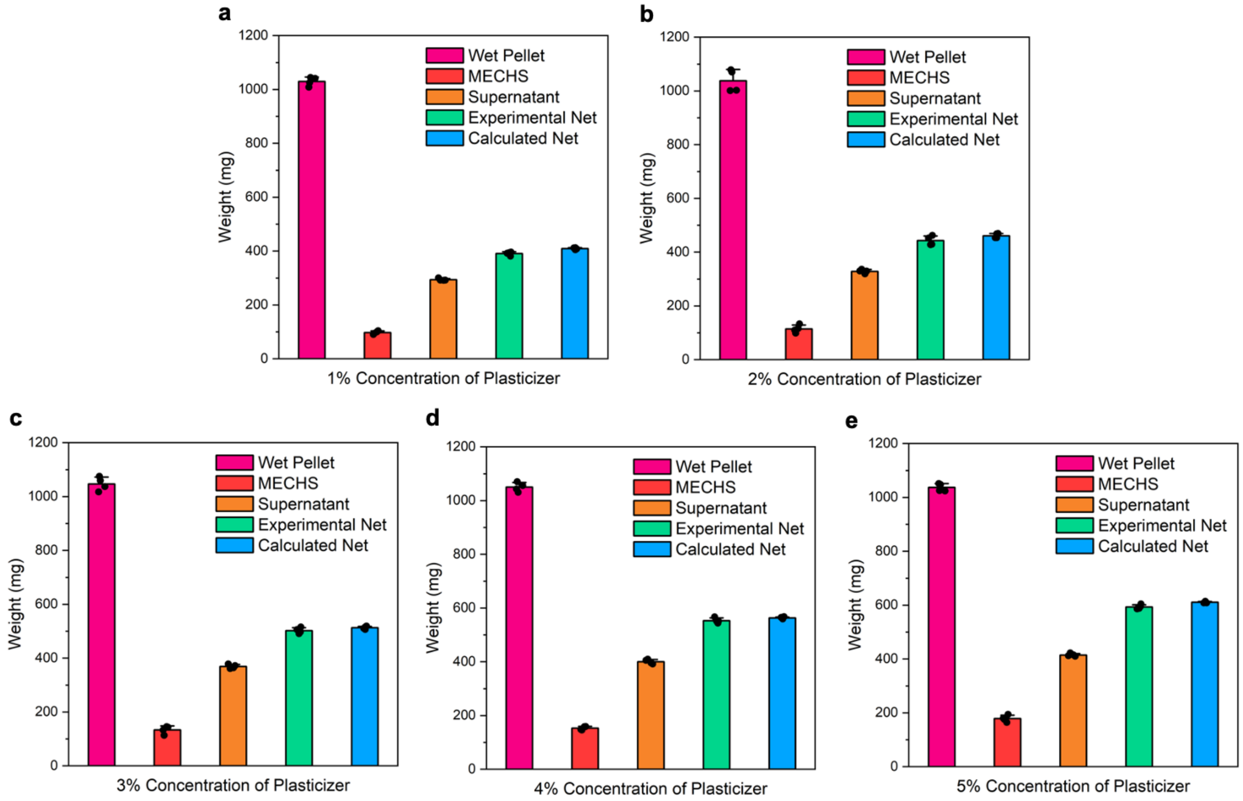


**Figure S15: Weight analysis of plasticizer treated MECHS.** Weights of wet pellet, MECHS film, dried supernatant, experimental net (MECHS and dried Supernatant) and calculated theoretical net (20.3% of wet pellet) for a) 1%, b) 2%, c) 3%, d) 4% and e) 5% of plasticizer. *n* = 4. Data represented as mean ± standard deviation.

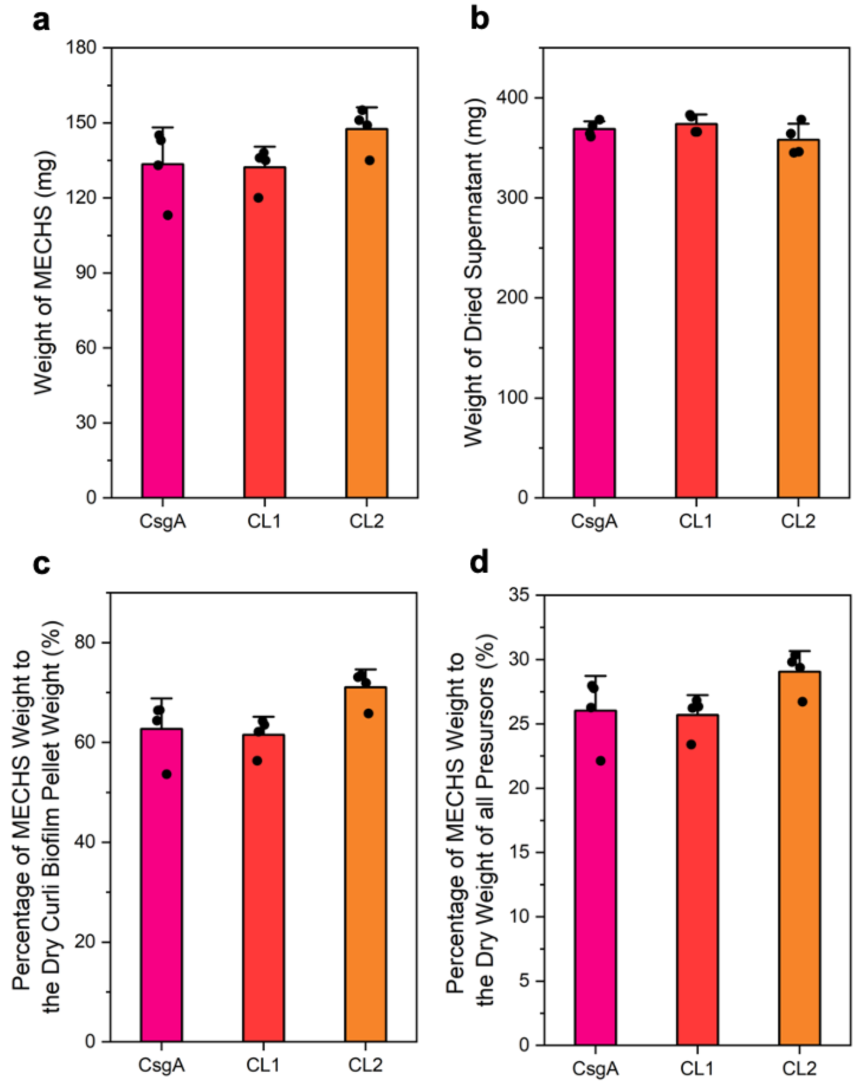


**Figure S16: Weight analysis of covalently crosslinked MECHS.** Weight of a) MECHS films and b) the dried supernatant of CsgA, CL1 and CL2. Percentage of MECHS weight to c) dry curli biofilm pellet weight (20.3% of wet pellet) and d) dry weight of all precursors for CsgA, CL1 and CL2. All precursors correspond to weights of cell pellet and that of 3% gelator and 1-5% plasticizer. *n* = 4. Data represented as mean ± standard deviation.

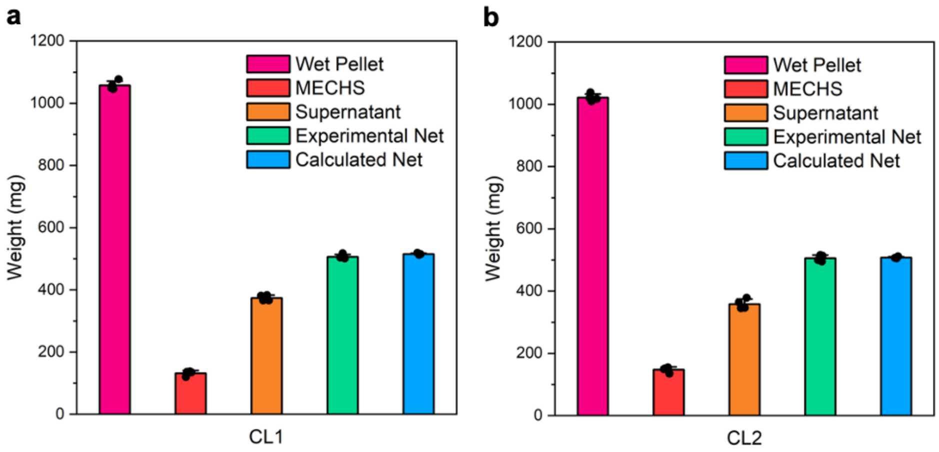


**Figure S17: Weight analysis of covalently crosslinked MECHS.** Weights of wet pellet, MECHS, dried supernatant, experimental net (MECHS and dried Supernatant) and calculated theoretical net (20.3% of wet pellet) for a) CL1 and b) CL2. $n$ = 4. Data represented as mean ± standard deviation.

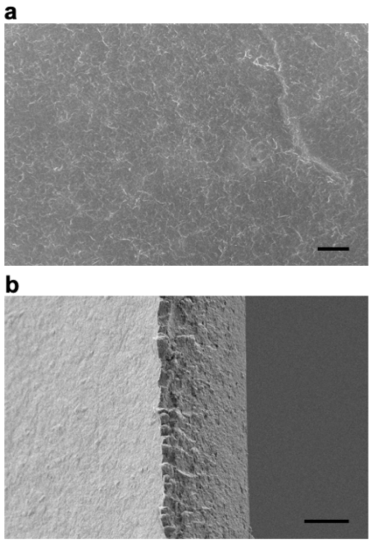

**Figure S18: Morphological analysis of MECHS.** FESEM image of a) top view and b) side view of MECHS film obtained from 3% gelator. Scale bar a) 20 μm and b) 10 μm.

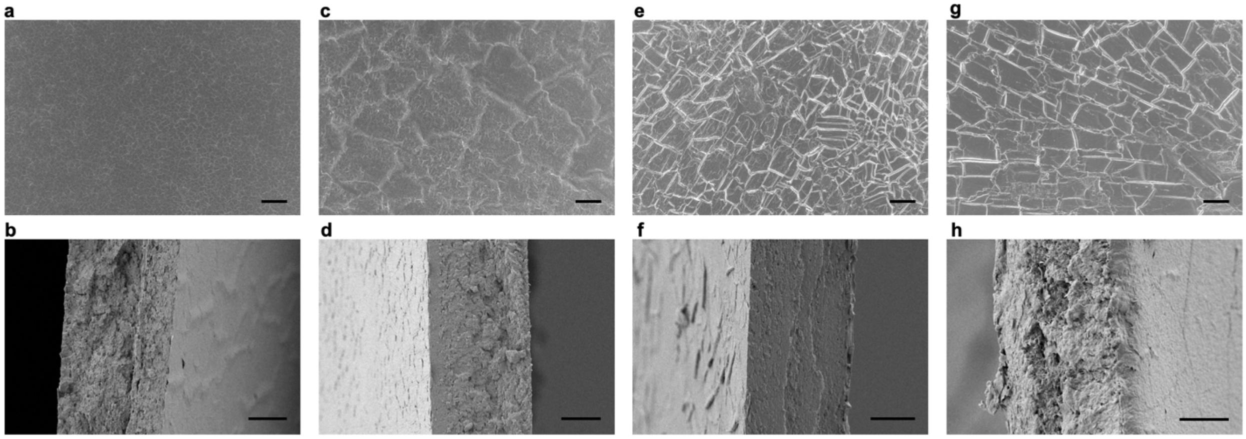


**Figure S19: Morphological analysis of MECHS.** FESEM image of a,c,e,g) top view and b,d,f,h) side view of MECHS film obtained from a,b) 1%, c,d) 2%, e,f) 4% and g,h) 5% plasticizer. Scale bar Top Row: 20 μm and Bottom Row: 10 μm.

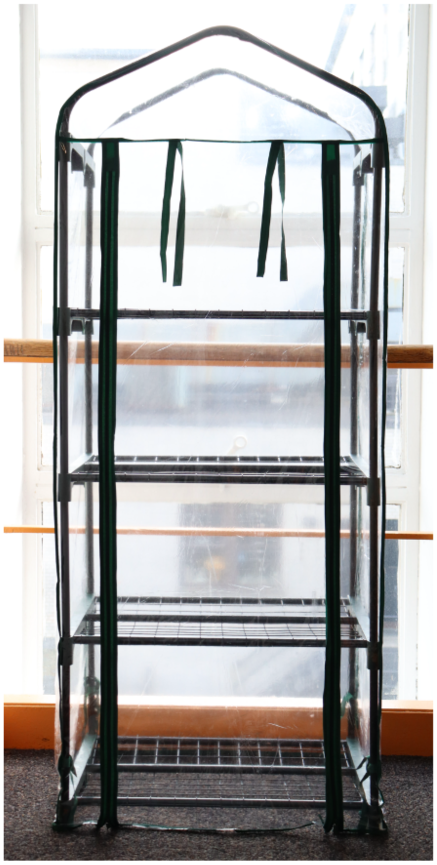

**Figure S20: Greenhouse for biodegradation experiment.** Photograph of a mini greenhouse setup utilized for testing the compostability of MECHS.

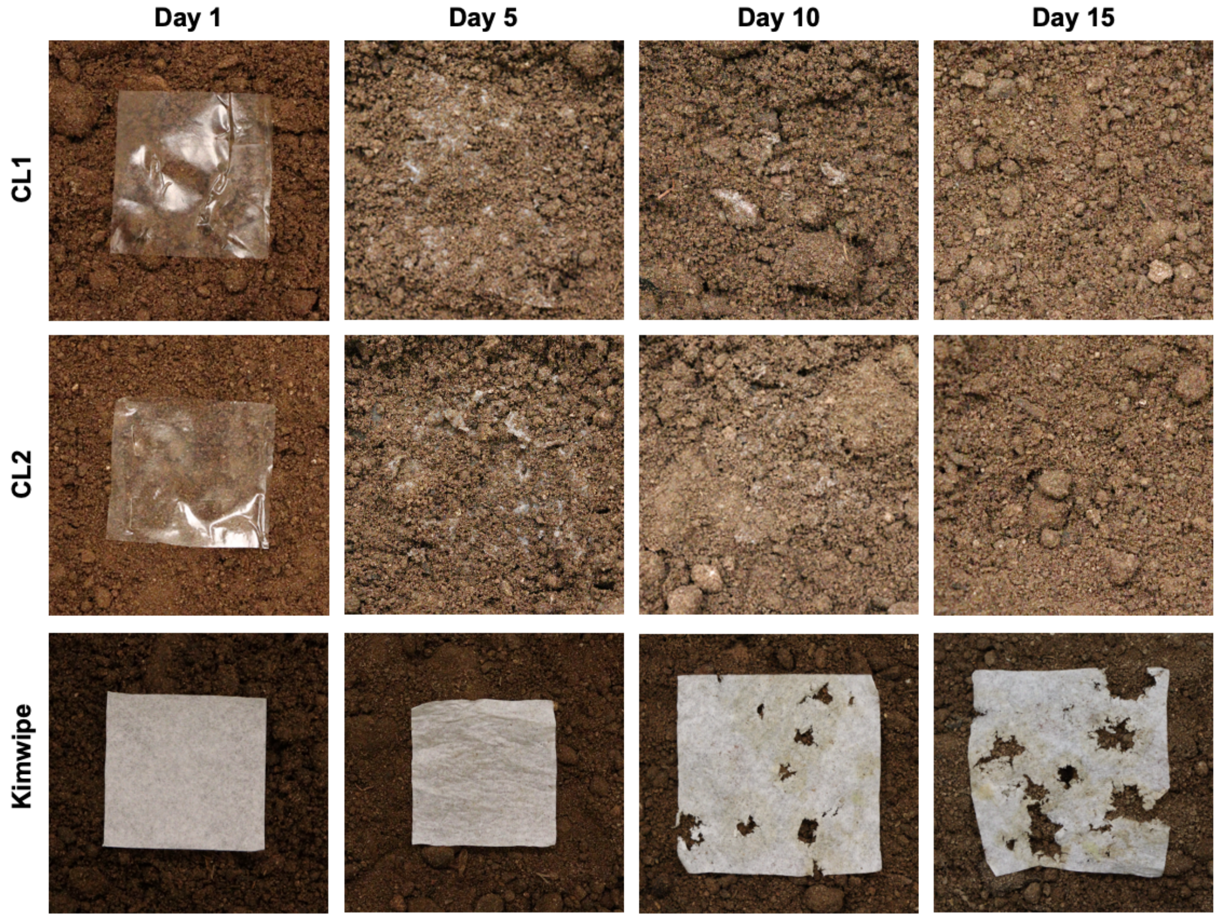


**Figure S21: Compostability of MECHS and Kimwipe in a fresh fishnure.** Photographs show the biodegradation of CL1, CL2 and Kimwipe in a fresh fishnure. The lateral dimensions of the samples were 5 cm by 5 cm.

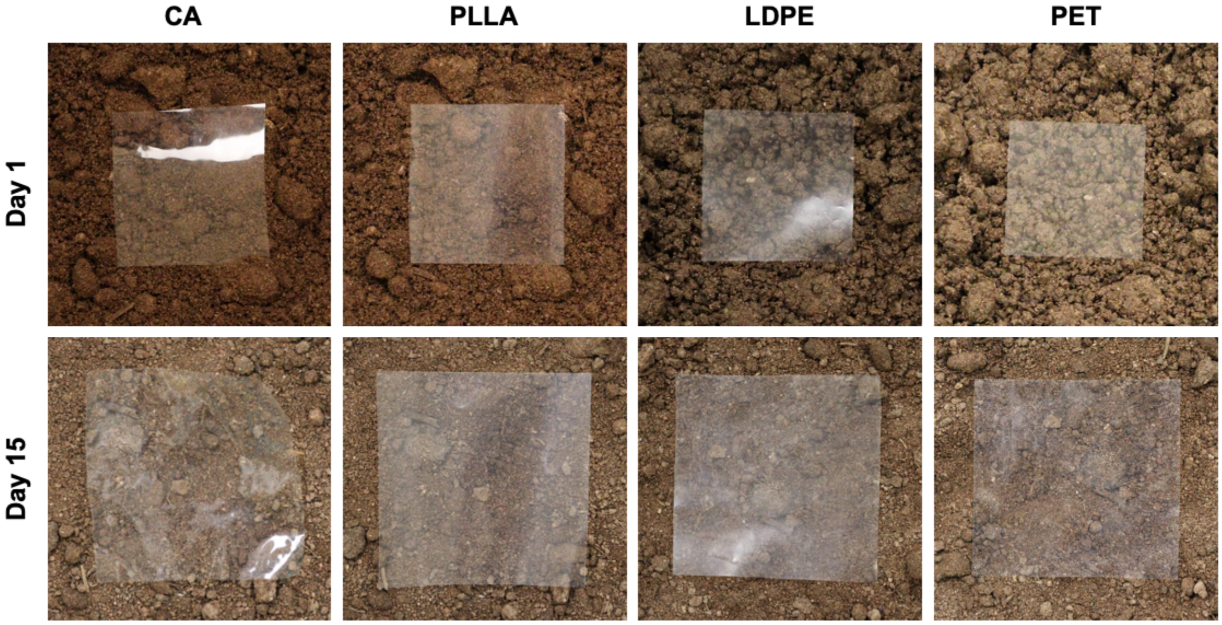


**Figure S22: Compostability of Bioplastics and Plastics in a fresh fishnure.** Photographs show the biodegradation of cellulose acetate (CA), poly-L-lactic acid (PLLA), low density polyethylene (LDPE) and polyethylene terephthalate (PET) in a fresh fishnure. The lateral dimensions of the samples were 5 cm by 5 cm.

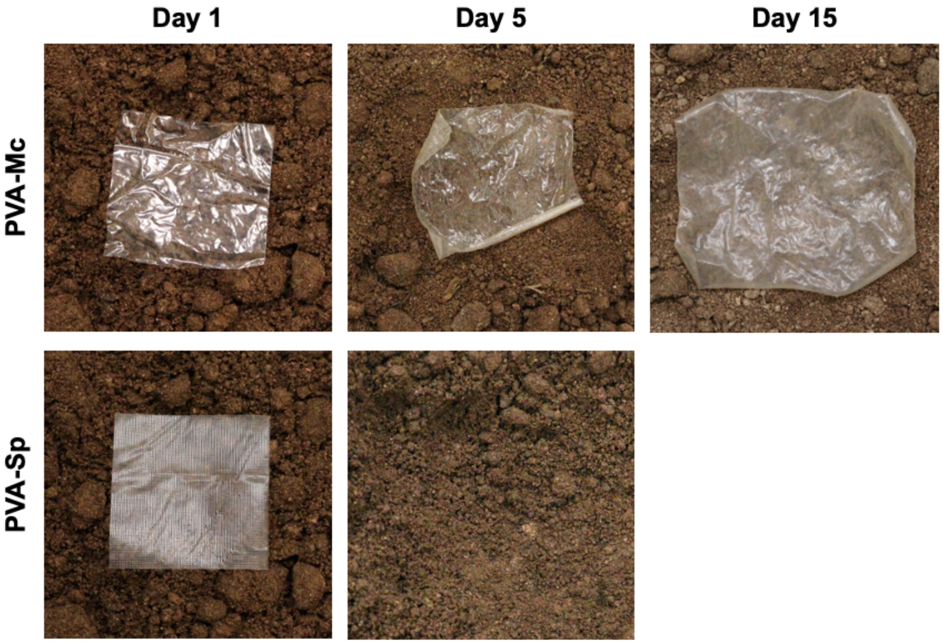


**Figure S23: Compostability of Polyvinyl alcohol in a fresh fishnure.** Photographs show the films of Polyvinyl alcohol - Mckesson (PVA-Mc) and Polyvinyl alcohol - Superpunch (PVA-Sp) in a fresh fishnure. The lateral dimensions of the samples were 5 cm by 5 cm.

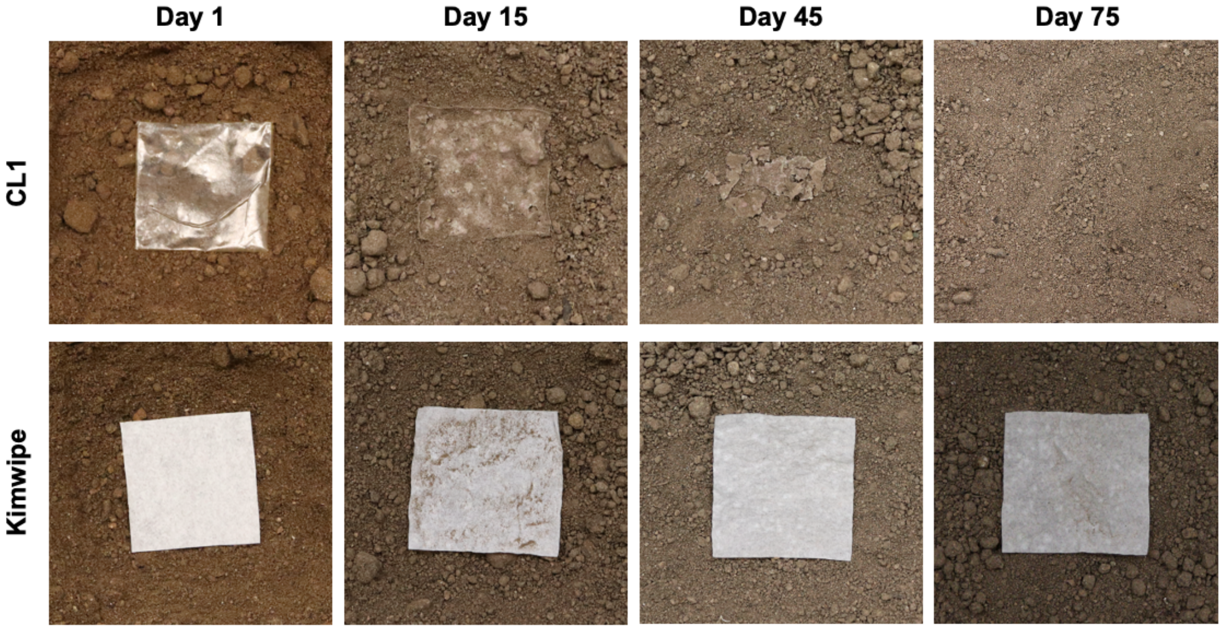


**Figure S24: Compostability of MECHS and Kimwipe in a dry fishnure.** Photographs show the biodegradation of CL1 and Kimwipe in a dry fishnure. The lateral dimensions of the samples were 5 cm by 5 cm.

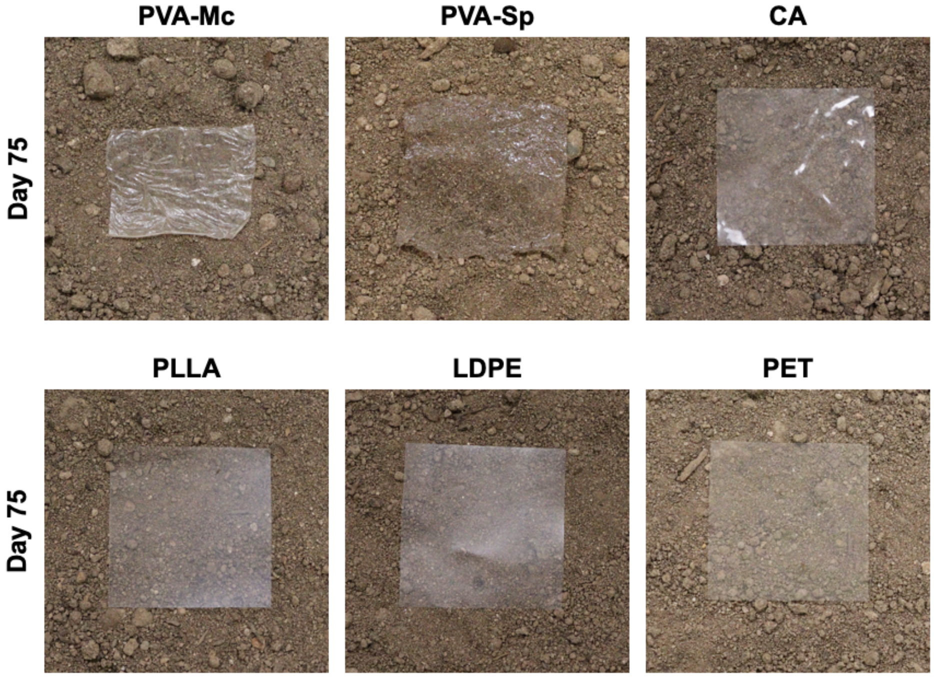


**Figure S25: Compostability of Bioplastics and Plastics in a dry fishnure.** Photographs show the biodegradation of Polyvinyl alcohol - Mckesson (PVA-Mc) and Polyvinyl alcohol - Superpunch (PVA-Sp), cellulose acetate (CA), poly-L-lactic acid (PLLA), low density polyethylene (LDPE) and polyethylene terephthalate (PET) in a dry fishnure. The lateral dimensions of the samples were 5 cm by 5 cm.

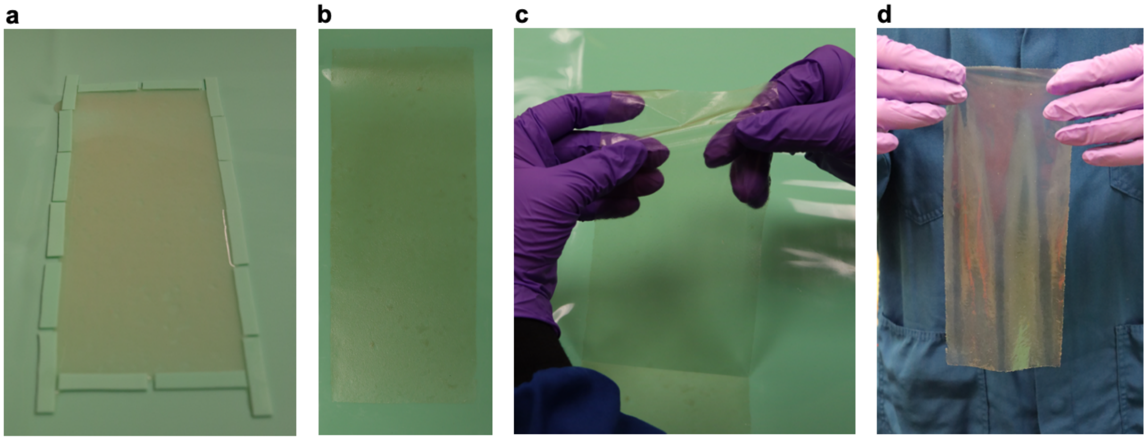


**Figure S26: Biofabrication of large MECHS prototype.** Photographs show the a) casting of curli biomass on to a silicone mold, b) ambient dried MECHS, c) peeling of MECHS film from the silicone mold and d) free-standing flexible MECHS film of lateral dimension 10 cm by 25 cm.

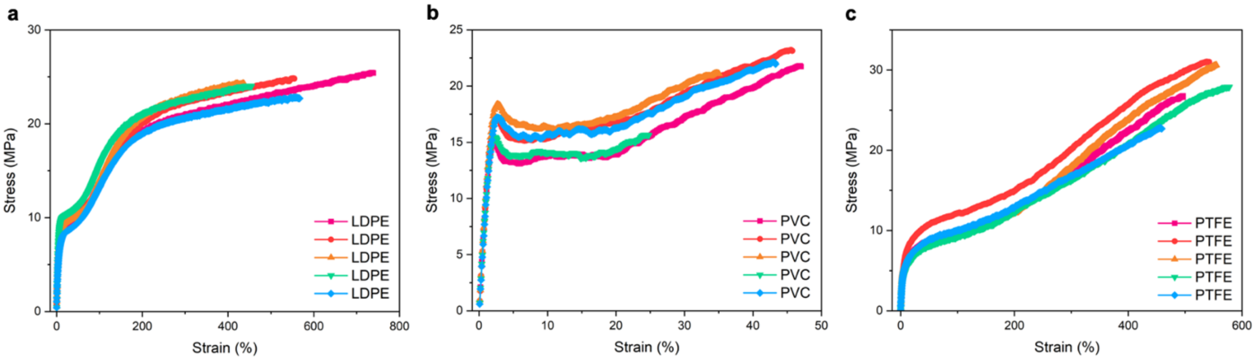


**Figure S27: Tensile tests.** Stress strain curves of a) LDPE, b) PVC and c) PTFE.

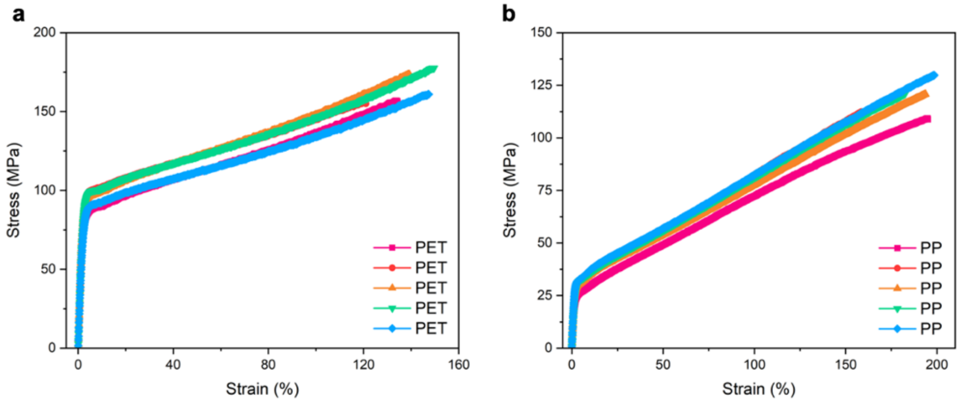


**Figure S28: Tensile tests.** Stress strain curves of a) PET and b) PP.

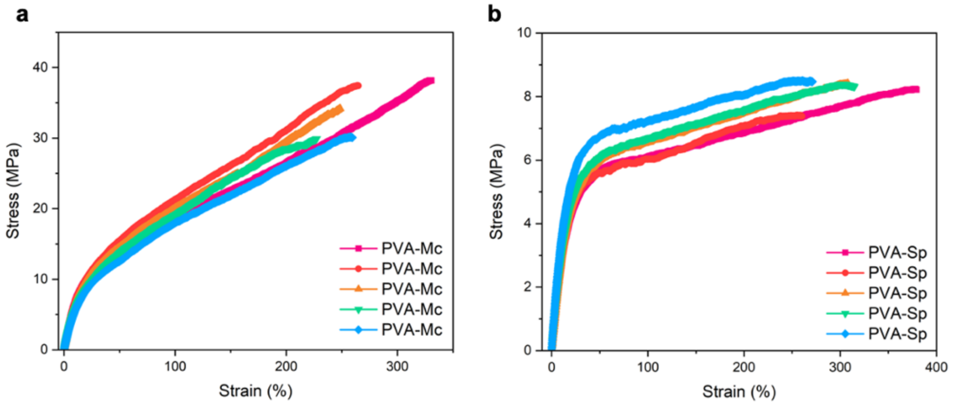


**Figure S29: Tensile tests.** Stress strain curves of a) PVA-McKesson and b) PVA-Superpunch.

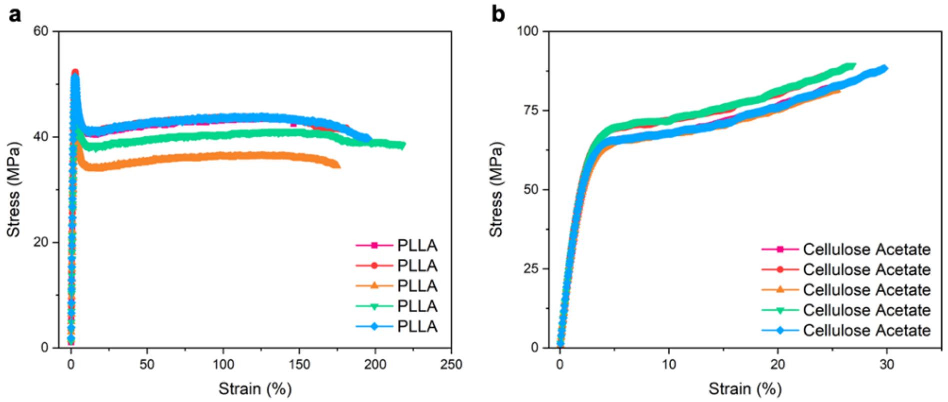


**Figure S30: Tensile tests.** Stress strain curves of a) PLLA and b) Cellulose Acetate.

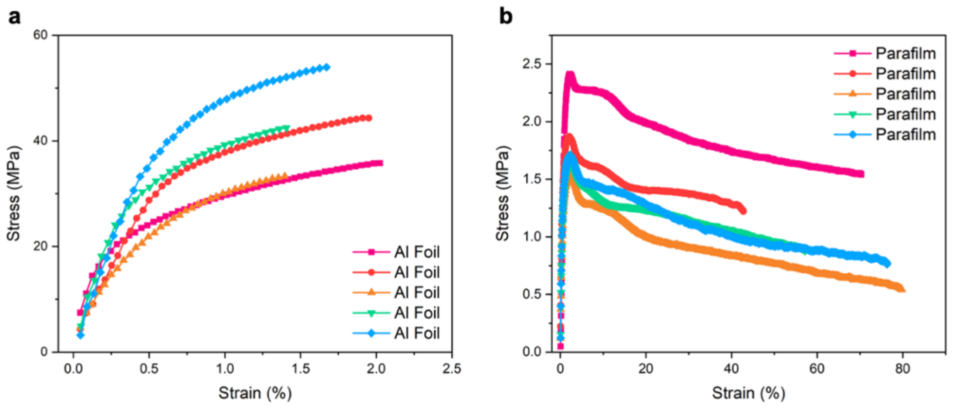


**Figure S31: Tensile tests.** Stress strain curves of a) Aluminum foil and b) parafilm.

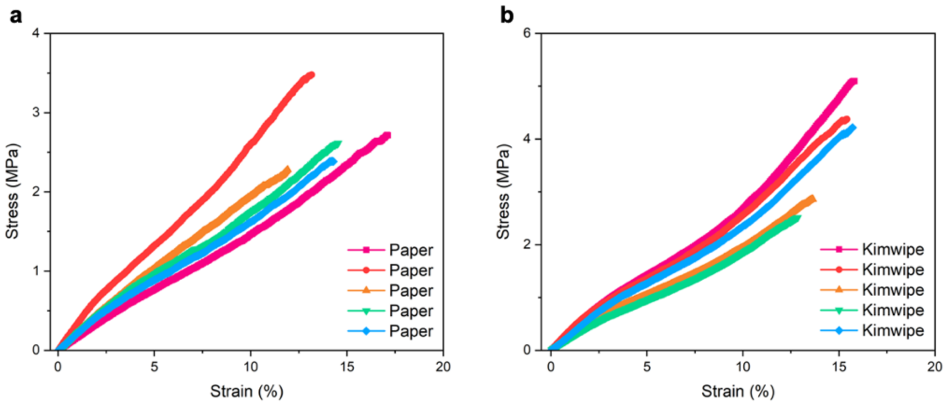


**Figure S32: Tensile tests.** Stress strain curves of a) Toilet paper and b) Kimwipe.

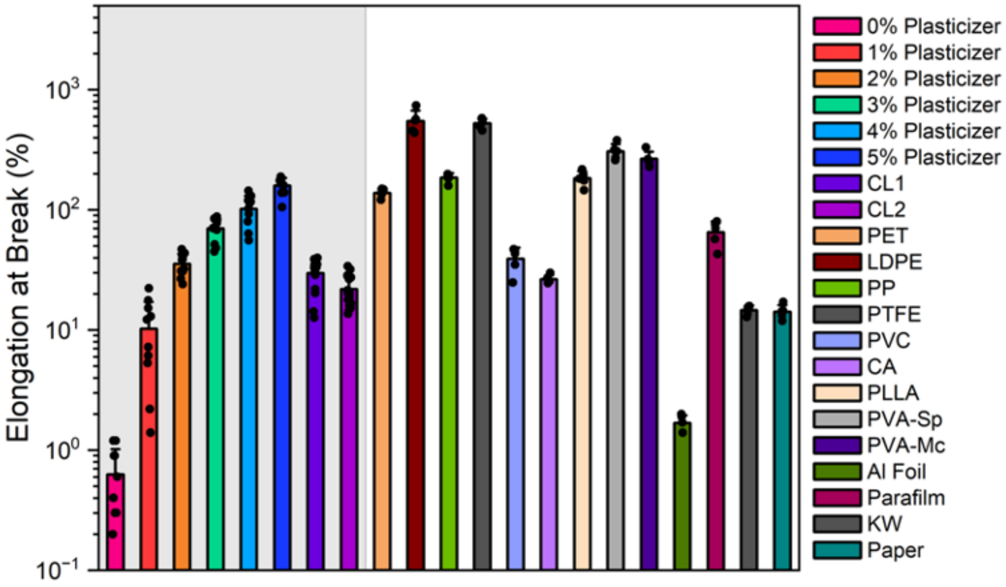


**Figure S33: Tensile tests.** Plot shows the elongation at break for MECHS, various synthetic materials and biomaterials.

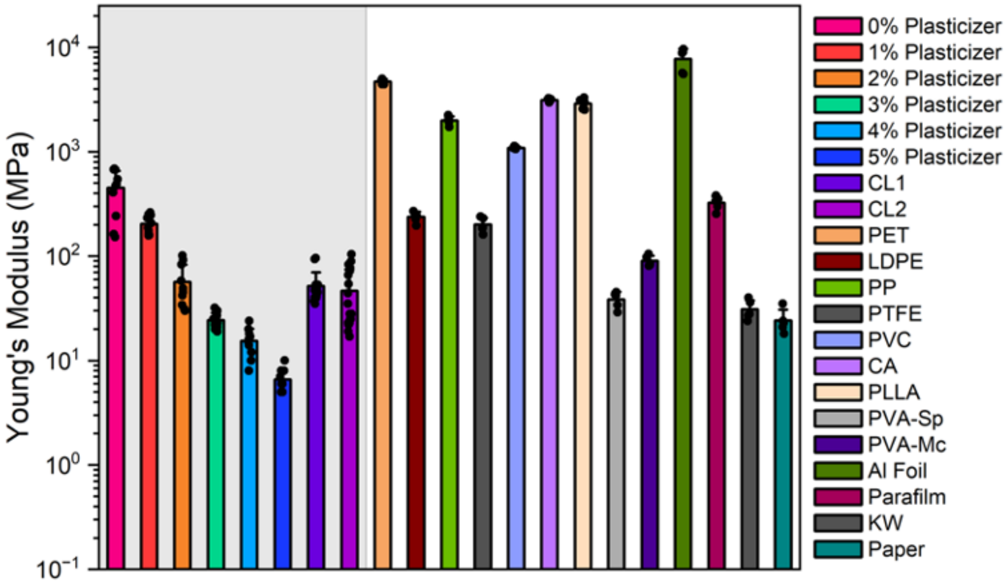


**Figure S34: Tensile tests.** Plot shows the Young's modulus for MECHS, various synthetic materials and biomaterials.

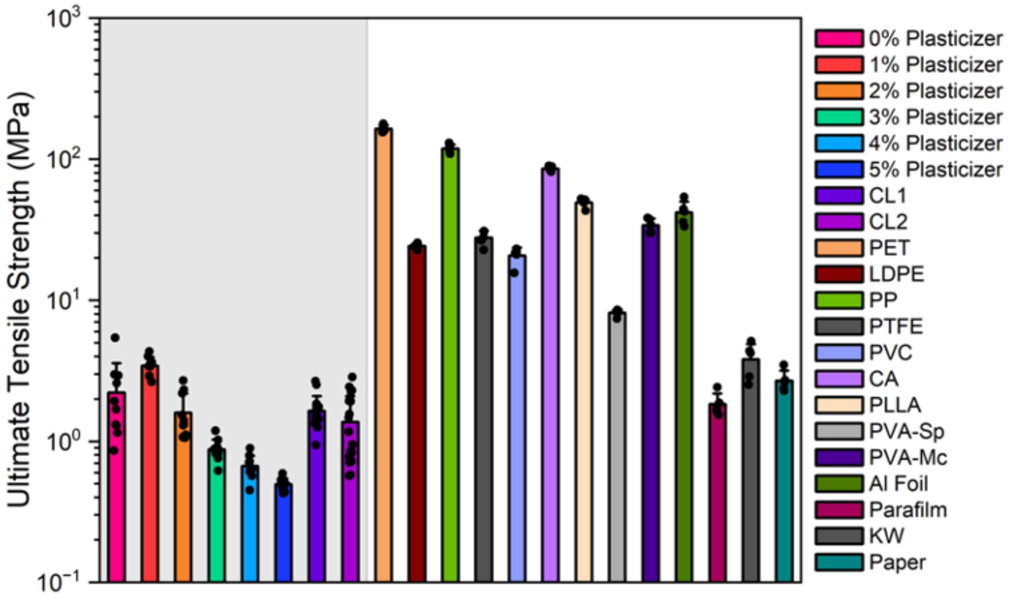


**Figure S35: Tensile tests.** Plot shows the ultimate tensile strength for MECHS, various synthetic materials and biomaterials.
